# Genome-wide Studies Reveal the Essential and Opposite Roles of ARID1A in Controlling Human Cardiogenesis and Neurogenesis from Pluripotent Stem Cells

**DOI:** 10.1101/2020.05.07.083113

**Authors:** Juli Liu, Sheng Liu, Hongyu Gao, Lei Han, Xiaona Chu, Yi Sheng, Yue Wang, Weinian Shou, Yunlong Liu, Jun Wan, Lei Yang

**Affiliations:** Department of Pediatrics, Indiana University School of Medicine, Herman B Wells Center for Pediatric Research, 1044 W Walnut Street, R4 272, Indianapolis, IN 46202, USA; Department of Medical and Molecular Genetics, Indiana University School of Medicine, Indianapolis, IN 46202, USA; Department of Obstetrics, Gynecology & Reproductive Sciences, University of Pittsburgh, Magee-Women’s Research Institute, Pittsburgh, PA 15213, USA; Center for Computational Biology and Bioinformatics, Indiana University School of Medicine, Indianapolis, IN 46202, USA; Department of BioHealth Informatics, Indiana University School of Informatics and Computing, Indiana University – Purdue University Indianapolis, Indianapolis, IN 46202, USA

**Keywords:** SWI/SNF, Chromatin Remodeling, ARID1A, REST, Cardiogenesis, Neurogenesis, Pluripotent stem cells

## Abstract

**Background:** Early human heart and brain development simultaneously occur during embryogenesis. Notably, in human newborns, congenital heart defects strongly associate with neurodevelopmental abnormalities, suggesting a common gene/complex underlying both cardiogenesis and neurogenesis. However, due to lack of *in vivo* studies, the molecular mechanisms that govern both early human heart and brain development remain elusive.

**Results:** Here, we report ARID1A, which is a DNA-binding-subunit of the SWI/SNF epigenetic complex, controls both neurogenesis and cardiogenesis from human embryonic stem cells (hESCs) via employing distinct mechanisms. Knockout of ARID1A (ARID1A^-/-^) led to spontaneous differentiation of neural cells together with globally enhanced expression of neurogenic genes in undifferentiated hESCs. Additionally, when compared with WT hESCs, cardiac differentiation from ARID1A^-/-^ hESCs was prominently suppressed, whereas neural differentiation was significantly promoted. Whole genome-wide scRNA-seq, ATAC-seq, and ChIP-seq analyses revealed that ARID1A was required to open chromatin accessibility on promoters of essential cardiogenic genes, and temporally associated with key cardiogenic transcriptional factors T and MEF2C during early cardiac development. However, during early neural development, transcription of most essential neurogenic genes was dependent on ARID1A, which could interact with a known neural restrictive silencer factor REST/NRSF.

**Conclusions:** We uncovered the opposite roles by ARID1A to govern both early cardiac and neural development from pluripotent stem cells. Global chromatin accessibility on cardiogenic genes is dependent on ARID1A, whereas transcriptional activity of neurogenic genes is under control by ARID1A, possibly through ARID1A-REST/NRSF interaction.

## Introduction

Human cardiogenesis concurs with neurogenesis and both have a similar process of morphogenesis including germ layer segregation, progenitor cell differentiation, cell fate specification, cell migration, left/right and dorsal/ventral patterning [1-3]. Developmental heart and brain have complex interactions at multiple levels. For example, neural crest cells (NCCs), capable of differentiation into peripheral neurons and glia, migrate from the pharyngeal arch arteries (PAA) and heart outflow tract (OFT) to the primitive heart, and give rise to smooth muscle, connective tissue, and great arteries of the heart [4]. Notably, newborns with congenital heart defects exhibited a high frequency of neurodevelopmental deficits [4, 5]. Previous studies revealed that epigenetic regulatory mechanisms, such as DNA methylation [6], histone modifications [7], and chromatin remodeling [8-11], played essential roles in heart and neural development in mammals. However, the molecular mechanisms driving both cardiogenesis and neurogenesis in human embryos remain unclear, which is critical for studying etiology of human congenital cardiac and neural defects.

The evolutionarily conserved ATP-dependent SWI/SNF complex is one of the largest chromatin remodeling complexes, consisting of ∼15 subunits, including SMARCA2 (also known as BRM) or SMARCA4 (also known as BRG1) as the ATPase catalytic subunit [12]. Several BRG1-associated factors (BAFs), such as ARID1A (Baf250a), have DNA binding capacity and assemble with either BRM or BRG1 to form a functional chromatin-remodeling complex [13]. BAFs also associate with other co-factors to determine the identity of a given SWI/SNF chromatin-remodeling complex and dictate where that complex will act [14]. As previously reported, ARID1A (Baf250a) binds promoter regions of transcription factors in a sequence-specific manner to drive SWI/SNF recruitment [15]. Loss of Ariad1a in mice results in embryonic lethality and developmental arrest at around day E6.5, showing lack of primitive streak or mesoderm formation [16]. A single amino acid mutation (Arid1aV^1068G/V1068G^), impaired Arid1a-DNA interactions [15] and resulted in neural defects including cranioschisis and neural tube defects, as well as cardiac defects including defective trabeculation, hypoplastic myocardial walls and ventricular septal malformation. Additionally, homozygous loss-of-ARID1a in mouse neural crest cells resulted in embryonic lethality at around E15, with prominent heart defects that included incomplete formation of the cardiac outflow tract septum and defective posterior pharyngeal arteries [17]. ARID subunits are among the most frequently mutated SWI/SNF subunits found in human disease [17]. Mutations in 4 different SWI/SNF subunits including ARID1A/B were identified in three congenital syndromes that include both neural and cardiac defects: Coffin-Siris syndrome (CSS), Nicolaides-Baraitser Syndrome (NCBRS), and ARID1B-related intellectual disability (ID) syndrome [13, 18-21]. Patients with those syndromes show severe intellectual deficits and cardiac defects, such as atrial/ventricular septal defects, patent ductus arteriosus (PDA), mitral and pulmonary atresia, aortic stenosis, and single right ventricle. These data indicate that malfunctional ARID1A may lead to defective formation of both heart and brain in humans. However, the detailed molecular mechanisms by which ARID1A controls human cardiogenesis and neurogenesis still remain elusive.

In this study, we investigate the roles of ARID1A in early human cardiac and neural development by using an in vitro human embryonic stem cell (hESC) model [22-25]. Expression of ARID1A is upregulated during early cardiac differentiation from hESCs. Surprisingly, knockout-of-ARID1A in hESCs (ARID1A^-/-^) led to spontaneous neural differentiation even under pluripotent stem cell culture conditions. Additionally, under conditions of targeted cardiac differentiation, ARID1A^-/-^ hESCs gave rise to robustly increased numbers of neural cells, including neural stem cells and neurons, whereas cardiac differentiation was significantly suppressed when compared with WT hESCs. ScRNA-seq revealed the cellular and transcriptional heterogeneities between ARID1A^-/-^ and WT hESCs. The Assay for Transposase-Accessible Chromatin using sequencing (ATAC-seq) in ARID1A^-/-^ and WT hESCs demonstrated changes in chromatin accessibility on ARID1A-occupied cardiogenic and neurogenic genes. ChIP-seq in ARID1A^-/-^ and WT hESCs revealed differences in genome-wide ARID1A occupancy on promoters of essential cardiogenic and neurogenic genes. When integrated, those genome-wide studies revealed loss-of-ARID1A globally reduced chromatin accessibility on essential cardiogenic genes, but did not uniformly affect chromatin accessibility on neurogenic genes. Furthermore, we performed Co-IP to show ARID1A interacted with key cardiac transcriptional factors T and MEF2C at the mesoderm and cardiac progenitor cell formation stages during human cardiogenesis. We identified that ARID1A recruited the transcriptional repressor element-1 (RE1) silencing transcription factor/neuron-restrictive silencer factor (REST/NRSF) to co-occupy the promoters of neurogenic genes, but not cardiogenic genes, for suppressing transcription of neurogenic genes. In summary, our findings reveal the distinct mechanisms by which ARID1A drives early human cardiogenesis and neurogenesis, primarily via chromatin remodeling and gene transcription regulation, respectively.

## Results

### Whole mRNA-seq predicts ARID1A in cardiac development from hESCs

Previously, we established a cardiovascular differentiation protocol from human embryonic stem cells (hESCs) by adding different combinations of growth factors including BMP4/bFGF/Activin A and VEGF/DKK1 at different stages [26]. Human ESC-derived multipotential cardiovascular progenitors (MCPs), cardiomyocytes (CMs), smooth muscle cells (SMs) and endothelial cells (ECs) were next enriched, followed with whole mRNA sequencing (Fig. 1a) [26, 27]. Gene Ontology (GO) biological process analysis by Metacore found the upregulated genes (MCP vs. hESCs) were significant enriched into cardiac development (Fig. 1b) and chromatin organization events (Fig. 1c), suggesting that chromatin organization/remodeling mechanism(s) could be underlying early human cardiac development. Metacore gene interaction network analysis of the upregulated genes (MCP vs. hESCs) found subunits of SWI/SNF complex, BAF250A (ARID1A) and BAF53A, may regulate expression of GATA4, which is a key cardiogenic transcription factor (Fig. 1d). This implies the potential role of SWI/SNF chromatin-remodeling complex in early human heart development. Next, we profiled expressions of essential SWI/SNF components in above five cell lineages by whole mRNA-seq (Fig. 1e). We found that expression level of ARID1A, a subunit of SWI/SNF complex with DNA binding capacity, significantly increased during hESCs differentiation to cardiac progenitors (Fig. 1e). QRT-PCR analysis confirmed the increased expression of ARID1A during cardiac differentiation of hESCs (Fig. 1f). Cardiac Troponin T (CTNT), which is a CM-specific marker gene, also showed increased expression during cardiac differentiation (Fig. 1f). Tissue-wide gene expression profiling demonstrated ARID1A in human adult heart tissue (Fig. S1a). Given that loss-of-Arid1a in mouse leads to severe developmental defects in both heart and neural tube [15], we decided to completely knock out ARID1A in H9 hESCs by using dual gRNAs mediated-CRISPR/Cas-9 technology (Figs. 1g, S1b). Two ARID1A^-/-^ hESC clones (#132 and #180) were identified by PCR (Figs. 1h, S1c). ARID1A mRNA (Fig. 1i) and protein (Fig. 1j) levels were undetectable in ARID1A^-/-^ clones when compared with WT hESCs. Large ARID1A^-/-^ hESC colonies displayed the same morphologies as WT hESCs. However, small cell clusters between large ARID1A^-/-^ colonies exhibited morphology of differentiated cell types (Figs. 1k, S1d). Those differentiated cell types were not observed in WT hESCs, indicating loss-of-ARID1A may induce sporadic differentiation of hESCs.

**Fig. 1.**
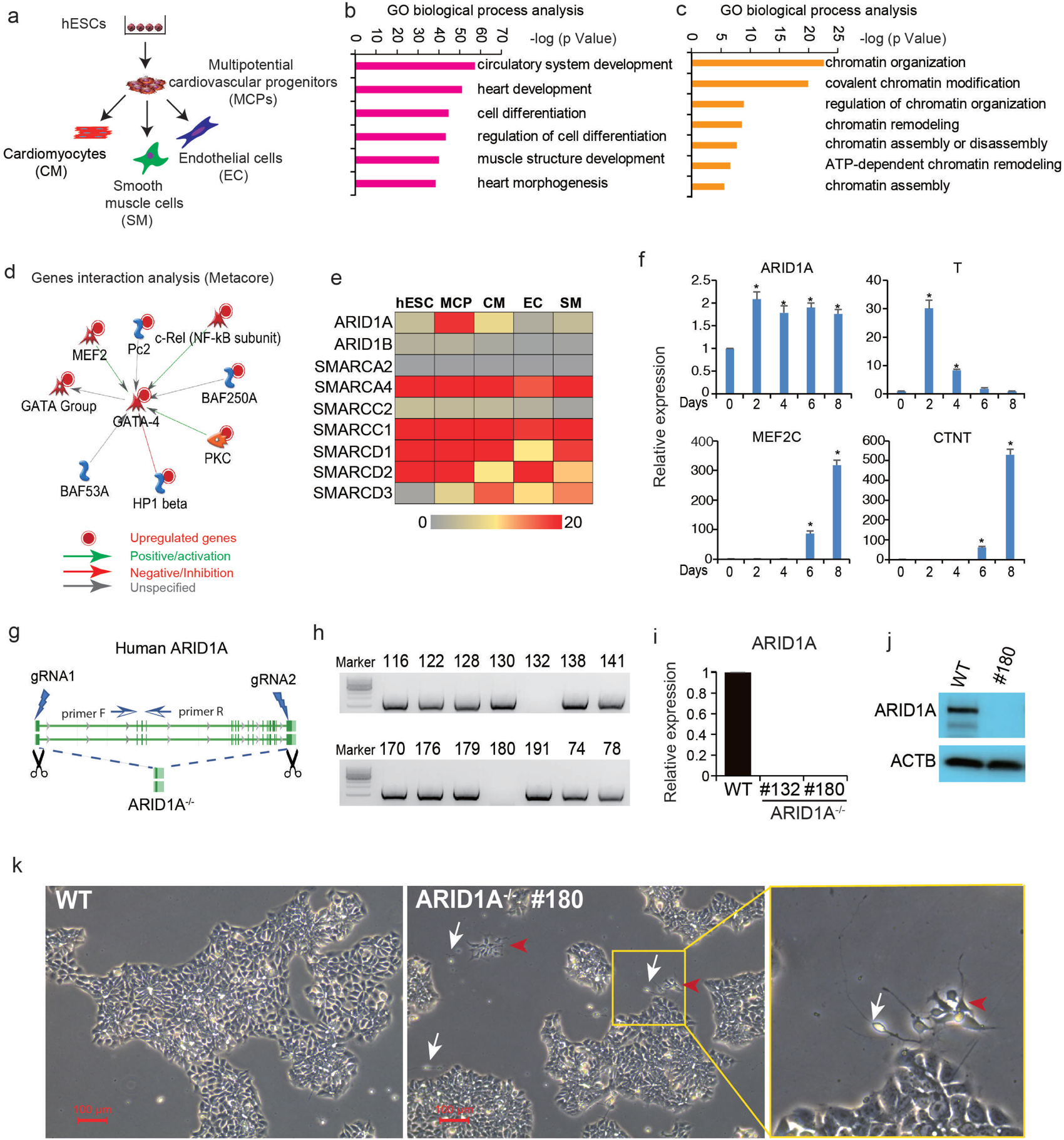
Loss-of-ARID1A induces spontaneous differentiation of hESCs. **a** Scheme for whole transcriptome sequencing of cardiovascular cell lineages derived from human embryonic stem cells (hESCs). **b-c** Gene Ontology (GO) biological process analysis of upregulated genes (MCP vs. hESCs) showing enriched heart development events (**b**) and epigenetic regulation events (**c**) during human cardiac development. **d** Interaction network analysis of SWI/SNF complex subunits and other cardiogenic genes by Metacore software. **e** Genes expression profiles of SWI/SNF subunits in different cardiovascular lineages by whole transcriptome sequencing. **f** Expression dynamics of ARID1A and key cardiac genes during cardiac differentiation of H9 hESCs. RNAs were collected every 2 days from day 0-day 10 of cardiac differentiation. **g** Strategy for completely knocking out human ARID1A in H9 hESCs using CRISPR/Cas9. **h** PCR screening for ARID1A^-/-^ hESC clones. **i** ARID1A mRNA expression detected by q-RT-PCR. All bars are shown as mean ± SD. n=3, \**p* < 0.05 (an unpaired two-tailed t-test with Welch’s correction). **j** ARID1A protein expression levels detected by Western Blot. **k** WT and ARID1A^-/-^ hESCs were cultured in mTesR medium. ARID1A^-/-^ hESCs displayed small clusters of differentiated cells (Red arrow heads and white arrows indicate two different cell types). All functional analysis and gene interaction network analyses were done by Metacore (Clarivate Analytics).

### Single cell RNA sequencing (scRNA-seq) reveals loss-of-ARID1A induces spontaneous neural differentiation

To determine the identity of differentiated ARID1A^-/-^ hESCs, single cell RNA sequencing (scRNA-seq) was performed (Fig. 2a). ARID1A mRNA was not detected in ARID1A^-/-^ hESCs, confirming the global ARID1A deficiency (Fig. 2b). Interestingly, expressions of neural stem cell markers (ZIC1, PAX6, SOX1) and neuron markers (MAP2, FABP7), were all significantly increased in ARID1A^-/-^ compared with WT hESCs (*p* < 2.4E-13, Fig. 2c). Consistently, the ratios of cells positively expressing ZIC1, PAX6, SOX1, MAP2 and FABP7 were all significantly higher in ARID1A^-/-^ hESCs than those in WT hESCs (*p* < 1.1E-14, Fig. 2d). In order to further compare the cellular heterogeneities, scRNA-seq data from both WT and ARID1A^-/-^ hESCs were integrated, followed with SEURAT, an R toolkit for single cell genomics [28], to perform dimension reduction and cell clustering. Totally 11 clusters were identified, as visualized by the t-distributed stochastic neighbor embedding (t-SNE) (Fig. 2e) [28, 29]. This revealed high heterogeneities of hESCs under pluripotency status. All clusters contained similar percentages of WT and ARID1A^-/-^ hESCs, except cluster 10. Particularly, over 90% cells in cluster 10 were from ARID1A^-/-^ hESCs (Fig. 2f). Cluster 10 had non-detectable level of ARID1A, but expressed neural stem cell markers ZIC1, SOX1 and neuron markers MAP2, FABP7, higher than any other clusters (Fig. 2g). In addition, neural-associated markers, such as PRTG, CDH6 and PTN, were highly expressed in cluster 10 (Fig. S2a). Gene Ontology (GO) (Fig. 2h), process networks and signaling pathways (Fig. S2b) analyses found that the upregulated genes (Table S1) in cluster 10 (ARID1A^-/-^ vs. WT) were significantly enriched in neural development events, such as nervous system development, neurogenesis, brain and head development. The feature plots showed that cells positively expressing neuroectoderm/neural stem cell markers (ZIC1, PAX6, SOX1) and neuron markers (MAP2, FABP7, PRTG, CDH6 and PTN) were highly enriched in cluster 10 and globally distributed in other clusters of ARID1A^-/-^ hESCs (red plots, Figs. 2i, S2c). Loss-of-ARID1A only decreased expression of OCT4 and did not affect the expression levels of other pluripotency markers (Fig. S2d), and endoderm (Fig. S2e), mesoderm and cardiac marker genes (Figs. S2f-h). Finally, flow cytometry, qRT-PCR and immunostaining were performed to verify results from scRNA-seq. Increased percentages of PAX6^+^, MAP2^+^ and TUJ1^+^ (Figs. 2j-k, S3a-c) cells were detected in ARID1A^-/-^ than WT hESCs by antibody staining followed with flow cytometry. Expression levels of neural-associated markers, such as PAX6, SOX1, MAP2, ZIC1 and FABP7, were significantly upregulated in ARID1A^-/-^ hESCs than those in WT (Figs. 2l, S3d). PAX6^+^, SOX1^+^, MAP2^+^ (Fig. 2m) and TUJ1^+^ (Fig. S3e) cells were detected in ARID1A^-/-^ hESCs, but not in WT hESCs, by fluorescent immunostaining. These results demonstrate that loss-of-ARID1A in hESCs increases transcription of neural genes and induces spontaneous neural differentiation (Figs. 2i, S3f), indicating ARID1A plays an important role in controlling neurogenesis from hESCs.

**Fig. 2.**
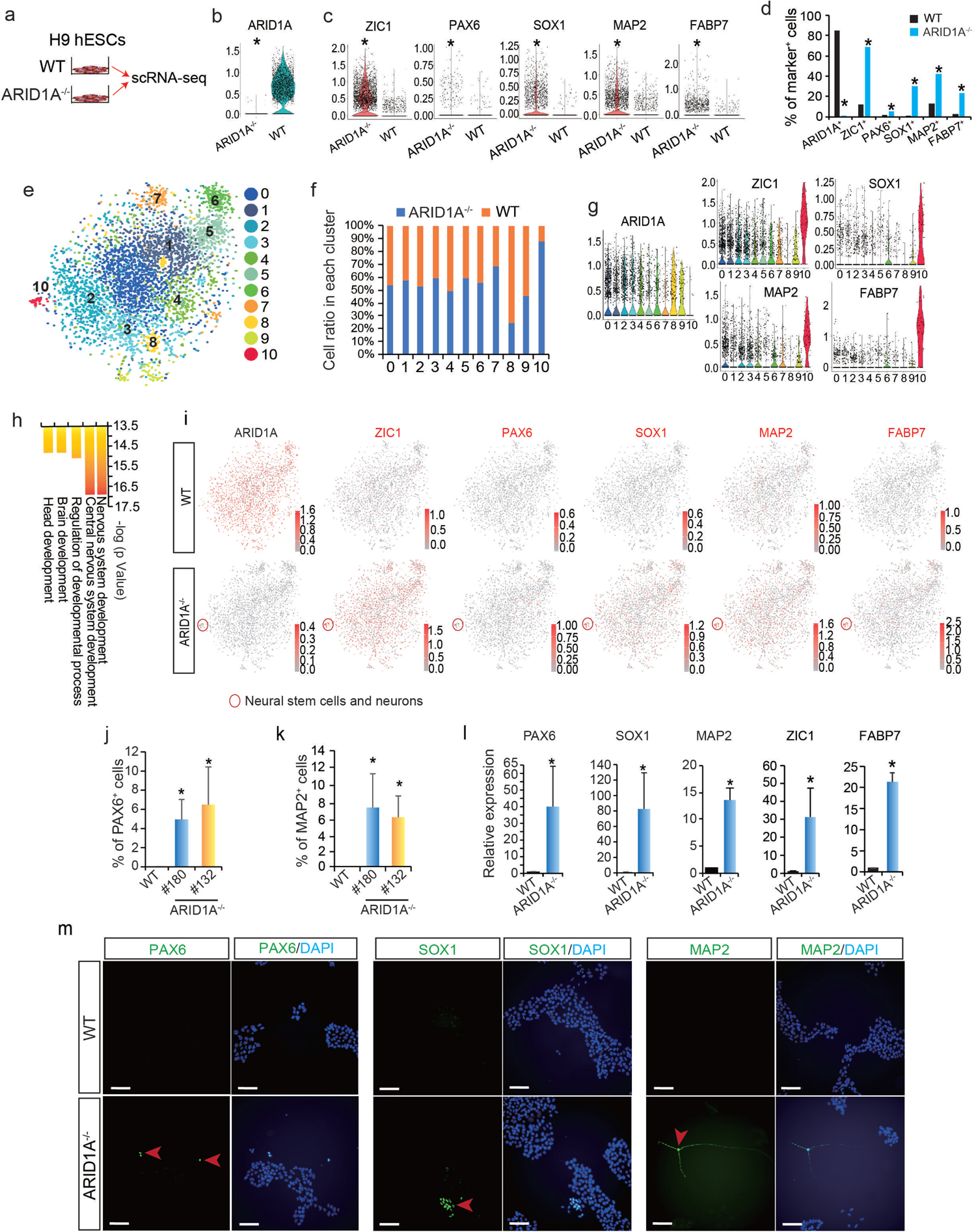
ScRNA-seq reveals loss-of-ARID1A induces spontaneous neural differentiation in hESCs. **a** Undifferentiated WT and ARID1A^-/-^ hESCs were collected for single cell RNA sequencing (scRNA-seq). **b** ARID1A expression in WT and ARID1A^-/-^ hESCs detected by scRNA-seq. \**p* < 2.4E-13 (Wilcoxon test). **c** Expression levels of neural stem cell markers (ZIC1, PAX6, SOX1) and neuron markers (MAP2, FABP7) in WT and ARID1A^-/-^ hESCs. \**p* < 2.4E-13 (Wilcoxon test). **d** Percentage of cells expressing ARID1A, neural stem cell markers (ZIC1, PAX6, SOX1) and neuron markers (MAP2, FABP7) in WT and ARID1A^-/-^ hESCs. \**p* < 1.1 E-14 (Fisher’s exact test). **e** Integrative analysis of scRNA-seq datasets from WT and ARID1A^-/-^ hESCs. Cell clusters were visualized with t-distributed stochastic neighbor embedding (t-SNE). **f** Ratios of WT and ARID1A^-/-^ hESCs in each cluster from integrative scRNA-seq data. **g** Violin plots of scRNA-seq data showing expression levels of ARID1A, neural stem cell markers (ZIC1, SOX1) and neuron markers (MAP2, FABP7) in each cluster. **h** GO biological process analysis for upregulated genes (ARID1A^-/-^ vs. WT) in cluster 10 by Metacore software. **i** Feature plots of scRNA-seq data showing the distribution of cells positively expressing ARID1A, neural stem cell markers (ZIC1, PAX6, SOX1) and neuron markers (MAP2, FABP7) in all clusters. Red circle indicates the cluster 10 containing neural stem cells and neurons solely derived from in ARID1A^-/-^ hESCs. **j-k** Flow cytometry data showing the percentage of PAX6^+^ (**j**) and MAP2^+^ (**k**) cells in WT and ARID1A^-/-^ hESCs. All bars are shown as mean ± SD. n=3, \**p* < 0.05 (KO versus WT, an unpaired two-tailed t-test with Welch’s correction). **l** Expression levels of neural-associated markers analyzed by qRT-PCR. All bars are shown as mean ± SD. n=3, \**p* < 0.05 (KO versus WT, an unpaired two-tailed t-test with Welch’s correction). **m** Immunomicroscopy to detect neural stem cell markers (PAX6, SOX1) and neuron marker (MAP2) in WT and ARID1A^-/-^ hESCs. Scale bar, 100µm.

### ARID1A oppositely controls cardiac and neural development from hESCs

Next, we asked whether loss-of-ARID1A could affect cardiac and neural differentiation from hESCs. A monolayer differentiation method [30] was utilized to induce cardiac differentiation from WT and ARID1A^-/-^ hESCs (Fig. 3a) for 10 days (T10), followed with scRNA-seq to distinguish cellular and transcriptional heterogeneities. ScRNA-seq detected no ARID1A expression in ARID1A^-/-^ hESCs post cardiac differentiation (Fig. S4a). By conducting Gene Ontology (GO) (Fig. 3b) and Process network (Fig. S4b) analyses of all differentially expressed genes (ARID1A^-/-^ vs. WT) (Table S1), we found the upregulated genes were significantly enriched into neural commitment events, such as nervous system development, neurogenesis and neural differentiation signaling pathway, whereas the downregulated genes were significantly enriched into cardiac commitment events, including circulatory system development, heart development, cardiac development and cardiac myogenesis signaling pathways. For example, loss-of-ARID1A led to increased expression of essential neural marker genes (NR2F1, OTX2, PAX6, SOX1, SOX2, ZIC1 and MAP2) (Fig. 3c) and reduced expression of cardiac marker genes (HAND1, GATA4, ISL1, NKX2-5, TNNT2, MYH6 and MYH7) (Fig. 3d) at day 10 of differentiation. Next, we quantified the percentages of single cells, which positively expressed neural and cardiac markers. Ratios of SOX1^+^, SOX2^+^, PAX6^+^, NR2F1^+^, OTX2^+^ and MAP2^+^ cells were prominently higher in ARID1A^-/-^ cells than WT cells (*p* < 3.13E-163, Fig. 3e). Oppositely, ratios of cells expressing cardiac markers, such as HAND1/2, TBX5, GATA4/5/6, ISL1, NKX2-5, MYH6/7, TNNT2, TNNI3 and TNNC1, were significantly lower in differentiated ARID1A^-/-^ than WT hESCs (*p* < 1.4E-114, Fig. 3f). ScRNA-seq data from WT and ARID1A^-/-^ cells were integrated to generate 12 clusters, which were visualized with t-distributed stochastic neighbor embedding (t-SNE) (Figs. 3g, S4c) [28, 29]. Next, the ratios of ARID1A^-/-^ and WT cells in each cell cluster were analyzed (Fig. 3h). Approximately 90% of cells in clusters 0, 6, 7 and 10 and 60% of cells in cluster 4 were derived from WT hESCs, whereas about 80% of cells in clusters 1, 2, 3, and 8 and 60% of cells in cluster 5 were derived from ARID1A^-/-^ hESCs. Cells expressing CM-specific marker genes, such as MYH6, TNNT2 (Fig. 3i), NKX2-5, GATA4/6, TBX5, MEF2C, MYH7, TNNI3 and TNNC1 (Fig. S4d), were enriched in clusters 0, 4, 6. Neural stem cell/neuroepithelial cell marker genes, such as ZIC1 (Fig. 3i), NR2F1, NR6A1, PAX6, SOX2/11 and ZIC2 (Fig. S4e), were highly expressed in cells of clusters 1, 2 and 3. Additionally, cells expressing neuron marker genes MAP2 (Fig. 3i), RBFOX3, SYP and DCX (Fig. S4e) were enriched in clusters 5, 8. Average gene expression in each cell cluster (ARID1A^-/-^ vs. WT) (Figs. S4f-k) was consistent with the results of violin plots. We conducted GO biological process and signaling pathway analyses of upregulated genes in each cluster. Cardiac/neural development events and pathways were separately enriched into clusters (Fig. S5). Feature plots from integrative analysis further distinguished distribution patterning of cells, which expressed signature neural and cardiac genes, across all clusters (Figs. 3j-k). Overall, scRNA-seq analysis found knockout- of-ARID1A predominantly enhanced differentiation of neural cells including neural stem cell, neuroepithelial cell and mature neuron from hESCs, whereas differentiation of cardiac cells including cardiomyocyte, endothelial cell and fibroblast was significantly suppressed (Fig. 3l). Given that cardiomyocyte, endothelial cell and fibroblast were originated from mesodermal cells, these results imply that loss-of-ARID1A could repress expression of genes controlling mesoderm formation.

**Fig. 3.**
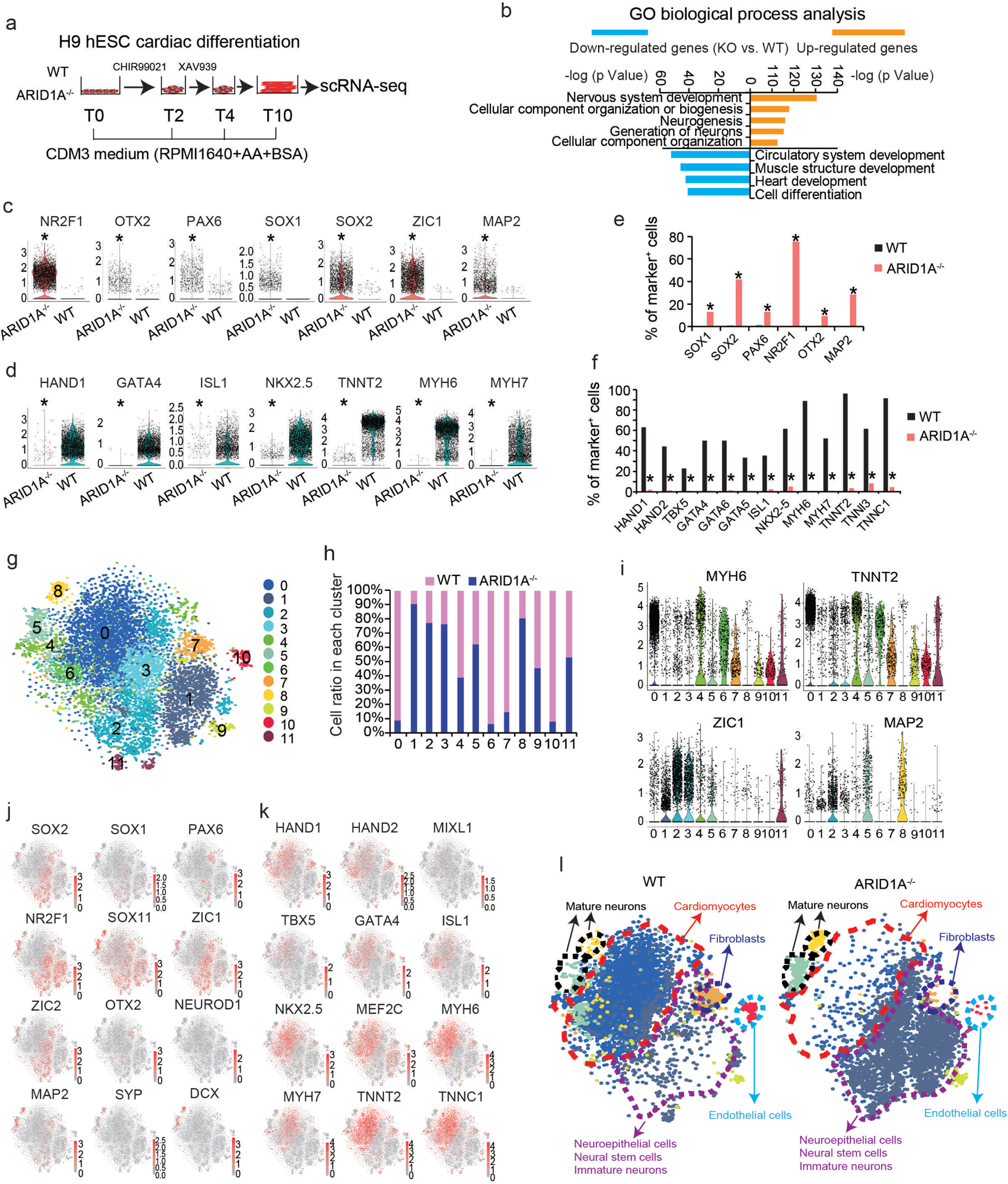
ScRNA-seq reveals loss-of-ARID1A represses cardiac but promotes neural differentiation from hESCs. **a** A chemically defined cardiac differentiation protocol to induce cardiomyocyte differentiation from hESCs, followed with scRNAseq. **b** GO biological process analysis of upregulated and downregulated genes (KO vs. WT differentiated cells) by Metacore software. **c-d** Violin plots showing expression of neural-associated markers (**c**) and cardiac-associated markers (**d**) in WT and ARID1A^-/-^ derived cells. \**p* < 8.32E-100 (Wilcoxon test). **e-f** Comparison of percentages of WT and ARID1A^-/-^ differentiated cells which positively express neural-associated markers (**e**) and cardiac-associated markers (**f**). * p<3.13E-163 (Fisher’s exact test) (**e**), * p<1.4E-114 (Fisher’s exact test) (**f**). **g** Integrative analysis of scRNA-seq datasets of differentiated WT and ARID1A^-/-^ hESCs. Cell clusters were visualized with t-distributed stochastic neighbor embedding (t-SNE). **h** Integrative analysis showing the ratios of WT and ARID1A^-/-^ cells in each cluster of differentiated cells. **i** Violin plots showing expressions of cardiac- and neural-associated markers in all clusters. **j-k** Feature plots showing the expressions and distributions of cells positively expressing neural-associated markers (**j**) and cardiac-associated markers (**k**) in the integrated view. **l** Defined cell types in the separated WT (left) and ARID1A^-/-^ (right) cell clusters. WT (left) clusters contain most cardiomyocytes and ARID1A^-/-^ (right) clusters contain most neural cells, indicating loss-of-ARID1A represses cardiac differentiation and promotes neural differentiation.

Next, after 10 days of cardiac differentiation (Fig. 3a), WT and ARID1A^-/-^ hESCs-derived cells were collected for FACS, qRT-PCR and immunostaining assessments. By performing FACS analysis, we found over 50% CTNT^+^ CMs from WT hESCs, whereas less than 20% CTNT^+^ CMs from ARID1A^-/-^ hESCs (Figs. 4a-b, Supplementary Videos: Differentiated WT and ARID1A KO hESCs). However, over 30% SOX1^+^ and TUJ1^+^ cells were derived from ARID1A^-/-^ hESCs, and less than 3% of those cells were derived from WT hESCs (Figs. 4c-d). QRT-PCR analysis found loss-of-ARID1A significantly repressed cardiac (Fig. 4e) but prominently increased neural marker gene expressions after differentiation (Fig. 4f). Immunomicroscopy detected robust CTNT^+^ CMs (Figs. 4g-h) and undetectable TUJ1+, MAP2+, SOX1+ and SOX2+ neural cells (Figs. 4g-j) derived from WT hESCs, whereas cells-derived from ARID1A^-/-^ hESCs gave opposite results (Figs. 4g-j). Next, a neural-specific differentiation protocol was performed to test whether loss-of-ARID1A can specifically affect neural commitment (Fig. S6a). We found ARID1A^-/-^ hESCs gave rise to significantly increased percentages of PAX6^+^ and SOX1^+^ cells than WT hESCs by FACS (Figs. S6b-c) and immunostaining (Figs. S6d-e). Expression levels of neural markers, such as PAX6, SOX1 and NEUROD1, increased post neural differentiation of ARID1A^-/-^ hESCs compared with WT hESCs (Fig. S6f). Enhanced neural differentiation was observed in ARID1A^-/-^ hESCs under both neural and cardiac differentiation conditions, indicating the master role of ARID1A in governing neural lineage specification from hESCs. Finally, since CRISPR/Cas9 completely ablated ARID1A expression in hESCs, we then asked whether dose of ARID1A could affect lineage commitment from hESCs. A single gRNA, targeting the transcriptional start site (TSS) of ARID1A with CRISPR/Cas9 (Fig. S6g), knocked down ARID1A expression to 60% of that in WT hESCs (Fig. S6h). After 10 days of cardiac differentiation, we found ARID1A^knockdown^ hESCs gave rise to half of CTNT^+^ CMs derived from WT hESCs (Fig. S6i). Additionally, shRNA-mediated knockdown of ARID1A (Fig. S6j) also confirmed that downregulation of ARID1A suppressed CM differentiation (Fig. S6k). However, compared with ARID1A^-/-^ hESCs, no sporadically differentiated neural cells were found in ARID1A^knockdown^ hESCs (Fig. S6l). Altogether, these results demonstrate that cardiac differentiation efficiency from hESCs is dependent on the dose of ARID1A, whereas spontaneous neural differentiation relies on the existence of ARID1A, implying different mechanisms by which ARID1A controls human cardiogenesis and neurogenesis.

**Fig. 4.**
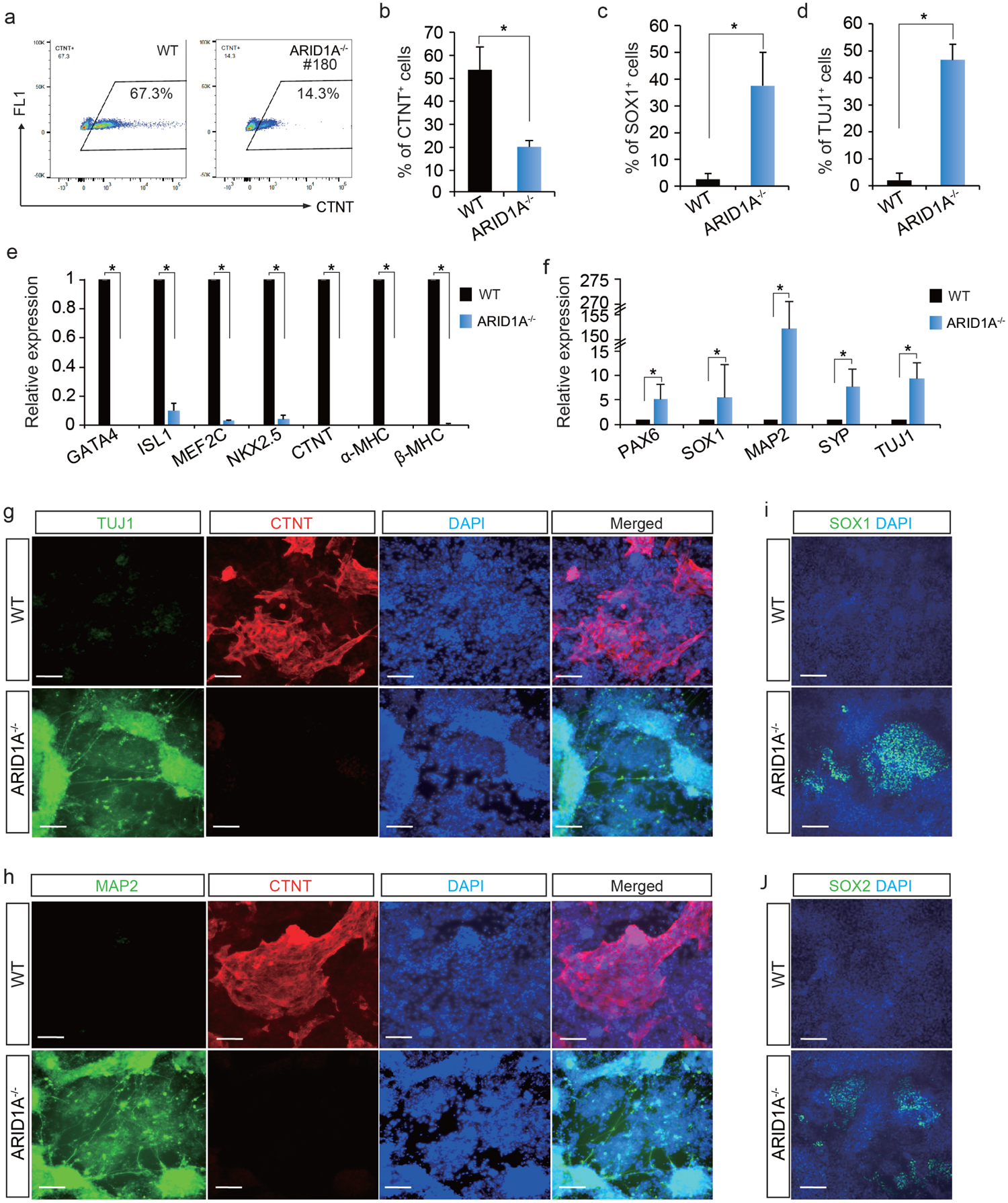
Characterization of WT and ARID1A^-/-^ hESC-derived cells. **a-b** Flow cytometry quantification of the ratios of CTNT^+^ CMs derived from WT and ARID1A^-/-^ cells after 10 days cardiac differentiation. All bars are shown as mean ± SD. n=3, \**p* < 0.05 (an unpaired two-tailed t-test). **c-d** Detecting ratios of SOX1^+^ (**c**) and TUJ1^+^ (**d**) cells in WT and ARID1A^-/-^ hESC-derived cells by immunostaining. All bars are shown as mean ± SD. n=3, \**p* < 0.05 (an unpaired two-tailed t-test). **e-f** Comparing the expression levels of cardiac markers (**e**) and neural markers (**f**) in WT and ARID1A^-/-^ hESC-derived cells by qRT-PCR. All bars are shown as mean ± SD. n=3, \**p* < 0.05 (KO versus WT for each gene, an unpaired two-tailed t-test). **g-h** Immunostaining of neuron markers TUJ1 (**g**), MAP2 (**h**) and cardiomyocyte marker CTNT (**g-h**) in WT and ARID1A^-/-^ hESC-derived cells after 10 days cardiac differentiation. Nuclei were labeled by DAPI. Scale bar, 100 µm. **i-j** Immunostaining of neural stem cell markers SOX1 (**i**) and SOX2 (**j**) in WT and ARID1A^-/-^ hESC-derived cells. Nuclei were labeled by DAPI. Scale bar, 100 µm.

### ATAC-seq reveals global alterations of chromatin remodeling in ARID1A^-/-^ hESCs

Given that SWI/SNF complex remodels chromatin to facilitate transcription of target genes, we next performed assay for transposase-accessible chromatin using sequencing (ATAC-seq) to investigate the impact of ARID1A on chromatin accessibility during cardiogenesis. HESCs at the day 4 (T4) of cardiac differentiation were collected for ATAC-seq (Fig. 5a). Global changes of chromatin accessibility, including increased (Fig. 5b), no change (Fig. 5g) and reduced (Fig. 5k), in ARID1A^-/-^ vs. WT hESCs were detected (Table S2). GO analysis found genes associated with differential chromatin accessibility changes have distinct biological functions. Genes with increased (Fig. 5b) or no-change (Fig. 5g) of chromatin accessibility (ARID1A^-/-^ vs. WT) were implicated into neural commitment events, such as nervous system development and neurogenesis (Figs. 5c-d, 5h, S7a). As shown in ATAC-seq (Fig. 5e) and qRT-PCR (Fig. 5f) data, loss-of-ARID1A increased both chromatin accessibility and expression levels of neurogenic genes, such as ZIC1, LHX5 and FABP7. However, chromatin accessibility on other neurogenic genes, such as PAX6, SOX1, FOXG1 (Fig. 5i), exhibited no obvious changes (ARID1A^-/-^ vs. WT), albeit their significantly increased expression levels (Fig. 5j). These results demonstrate that transcriptional activities, rather than chromatin accessibility, of critical neurogenic genes were dependent on ARID1A. Notably, genes with reduced chromatin accessibility (ARID1A^-/-^ vs. WT) (Fig. 5k) were mainly enriched into cardiac commitment events, such as circulatory, heart and cardiovascular system development (Figs. 5l, S7b). These genes included early mesoderm transcriptional factors T [31-37], cardiac transcription factors GATA4, ISL1, TBX3, DES, NKX2.5, TBX5 and TBX18, and sarcomeric genes CTNT, MYH7 (Figs. 5m-n). Additionally, during cardiac differentiation, expression levels of these cardiogenic genes were all significantly decreased in ARID1A^-/-^ cells compared with WT cells (Fig. 5o). Loss-of-ARID1A did not affect the expression of endoderm marker genes, such as SOX17 and FOXA2 (Fig. S7c). Altogether, these results indicate that loss-of-ARID1A globally reduces chromatin accessibility on essential cardiogenic genes, but not on neurogenic genes, whereas transcription of essential neurogenic genes is dependent on ARID1A.

**Fig. 5.**
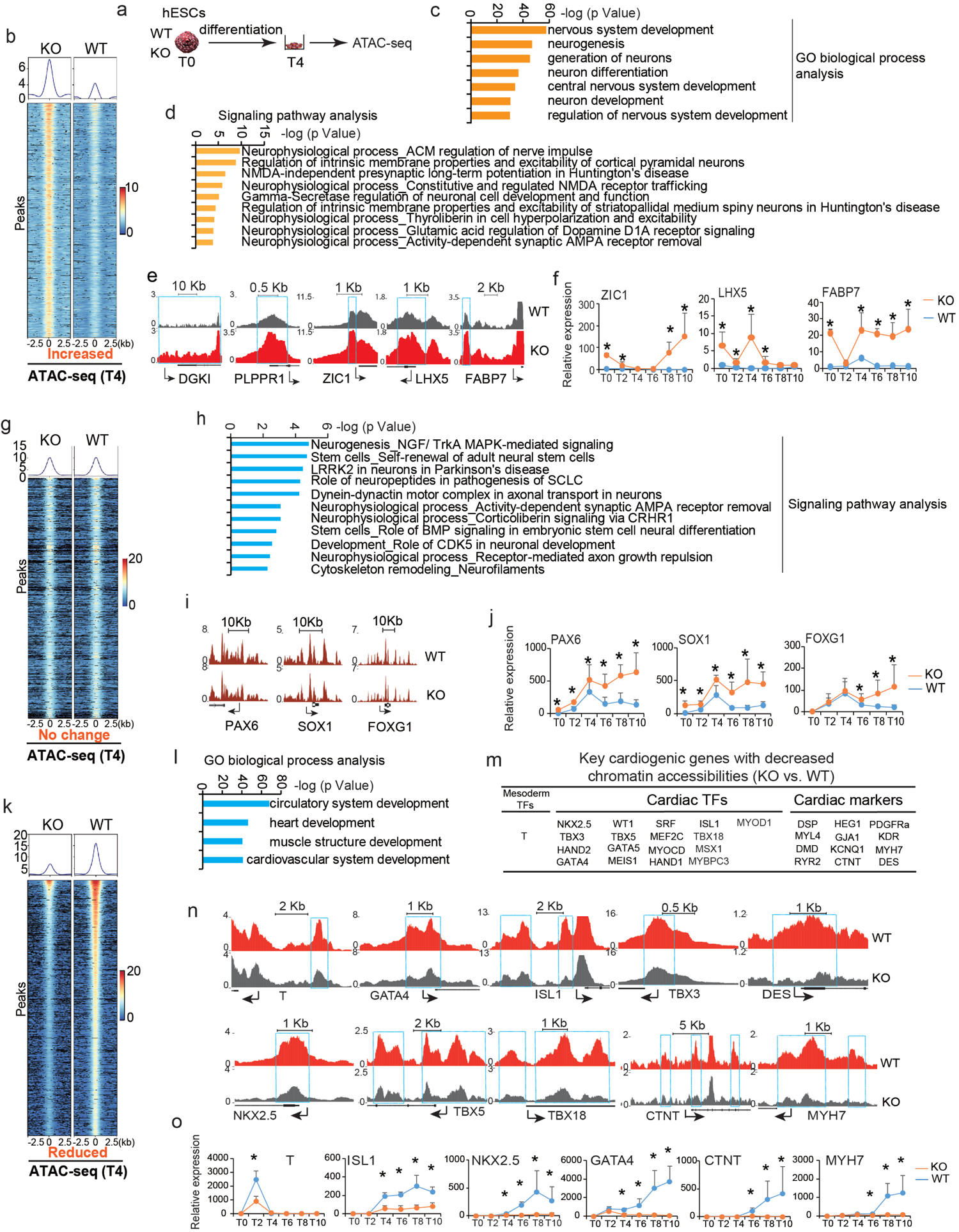
Loss-of-ARID1A leads to changes of chromatin accessibility. **a** Assay for Transposase Accessible Chromatin with high-throughput sequencing (ATAC-seq) was performed after 4 days differentiation of WT and ARID1A^-/-^ hESCs. **b** Heatmap and pileup of ATAC-seq signal for genes with increased chromatin accessibility (KO vs. WT). **c-d** GO biological process (**c**) and Signaling pathways enrichment (**d**) analysis of genes with increased chromatin accessibility (KO vs. WT). The analysis of GO biological process and signaling pathways were performed by Metacore software. **e** Genome views of neurogenic genes with increased chromatin accessibility (KO vs. WT) close to transcription start site (TSS). Arrows showed the transcriptional direction. **f** QRT-PCR detection of neurogenic gene expression profiles during cardiac differentiation of hESCs. **g** Heatmap and pileup of ATAC-seq signals for genes without significant changes on chromatin accessibility (KO vs. WT). **h** Signaling pathways enriched in genes with no-changed chromatin accessibility (KO vs. WT). The analysis of signaling pathways was performed by Metacore software. **i** Genome views of neurogenic genes without significant changes on chromatin accessibility (KO vs. WT). **j** QRT-PCR detection of neurogenic gene expression profiles during cardiac differentiation of hESCs. **k** Heatmap and pileup of ATAC-seq signals for genes with reduced chromatin accessibilities (KO vs. WT). **l** GO biological process analysis of genes with reduced chromatin accessibilities (KO vs. WT). The analysis of GO biological process was performed by Metacore software. **m** List of essential cardiogenic and cardiac functional genes showing decreased chromatin accessibility in KO cells (KO vs. WT). **n** Genome views of some important cardiogenic genes with decreased chromatin accessibility in KO cells compared to WT. Arrows showed the transcriptional direction. **o** QRT-PCR detection of cardiogenic gene expression profiles during cardiac differentiation of hESCs. All bars in RT-QPCR data are shown as mean ± SD. n=3, *p < 0.05 (KO vs. WT, an unpaired two-tailed t-test with Welch’s correction).

### ChIP-seq reveals genome-wide ARID1A occupancy on promoters of essential cardiogenic and neurogenic genes

Given the DNA binding capacity of ARID1A [15], chromatin immunoprecipitation assay with sequencing (ChIP-seq) was next carried out to define genome-wide ARID1A occupancy in hESCs (Fig. 6a). Approximately 61% of ARID1A binding sites located on regions within 10 kb from transcriptional start sites (TSS), 5’UTR, and gene bodies (Fig. 6b). Interestingly, GO analyses revealed that genes with ARID1A occupancies close to TSS (Fig. 6c, Table S3) were enriched into both neural and cardiac commitments, such as nervous system development, neurogenesis (Fig. 6d) and cardiac development/muscle contraction (Fig. 6e). For example, transcriptional factors for early mesodermal commitment, EOMES, and key cardiac transcriptional factors, HAND2, GATA4, GATA5, ISL1, TBX3, were all occupied by ARID1A on their promoter regions (Figs. 6f-g), which were confirmed by ChIP-qPCR (Fig. 6h). Both ChIP-seq and ChIP-qPCR also identified ARID1A-occupied neurogenic genes, such as PAX6, SOX1, FOXG1 and ZIC1 (Figs. 6i-k). Importantly, by comparing the ATAC-seq (Fig. 5m) and ARID1A ChIP-seq data (Fig. 6g), we found all ARID1A-occupied essential cardiogenic genes exhibited decreased chromatin accessibility during cardiac commitment from ARID1A^-/-^ vs. WT hESCs, indicating that chromatin accessibility of cardiogenic genes was dependent on ARID1A. Although ARID1A occupied promoters of many neurogenic genes (Fig. 6j), most of them, except several genes including ZIC1, LHX5 and FABP7 (Fig. 5e), did not exhibit significant alternations of chromatin accessibility under ARID1A deficiency. This demonstrates that ARID1A globally occupies neurogenic gene promoters to suppress transcription via a non-chromatin-remodeling mechanism. It also implies the existence of ARID1A-recruited co-factor(s) to repress neurogenic gene transcription.

**Fig. 6.**
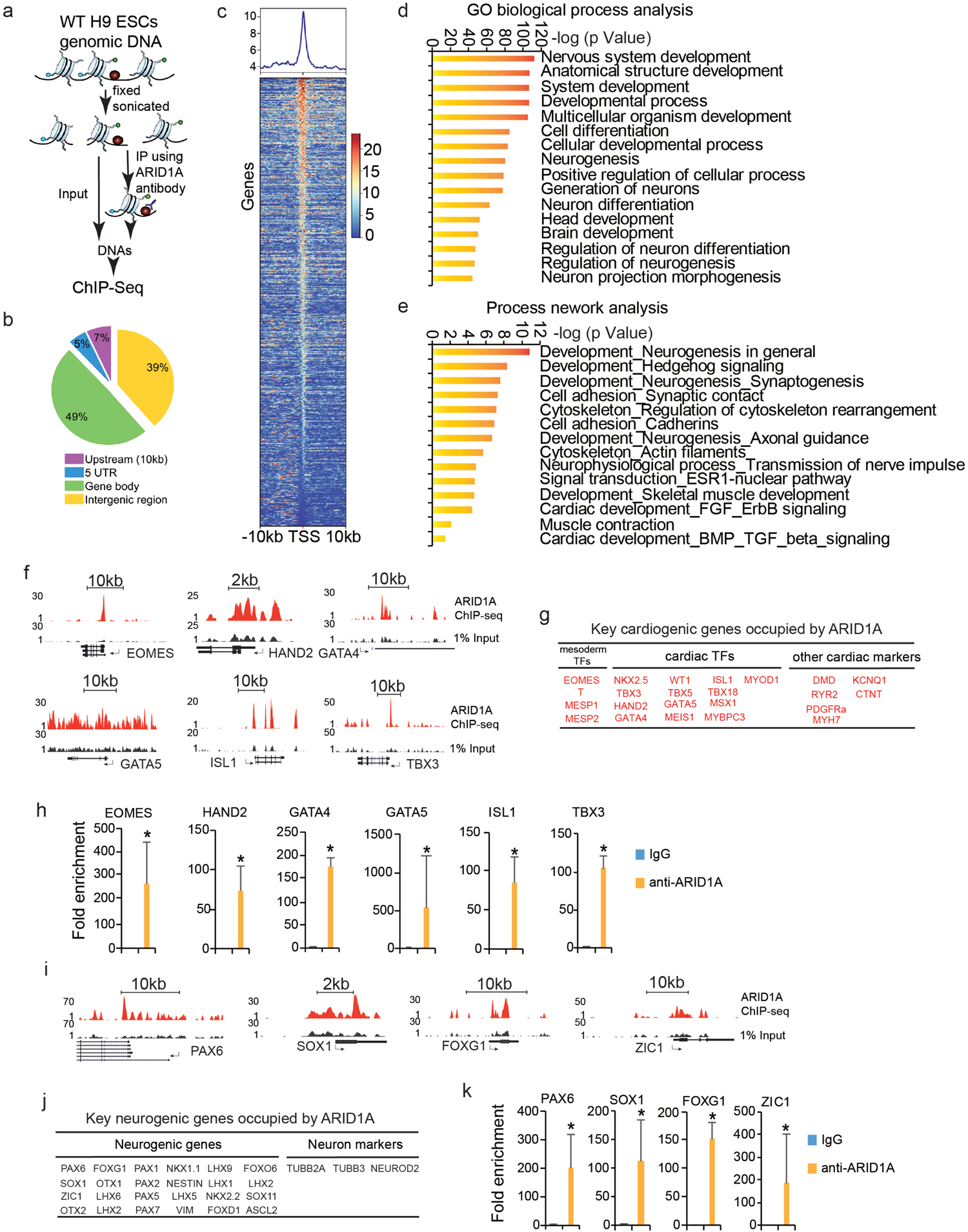
ChIP-seq reveals genome-wide ARID1A occupancies on promoters of cardiogenic and neurogenic genes. **a** Chromatin immunoprecipitation (ChIP) assays with sequencing (ChIP-Seq) was performed in WT H9 hESCs. **b** Distribution of ARID1A ChIP-seq-enriched peaks across human genome. **c** Relative enrichment levels of ARID1A on transcription start site (TSS) of coding genes in hESCs. Heatmaps were ranked by ARID1A enrichment. **d** GO biological process analysis of genes enriched by ARID1A ChIP-seq. The analysis of GO biological process was performed by Metacore software. **e** Process networks analysis of genes enriched by ARID1A ChIP-seq. The analysis of process networks was performed by Metacore software. **f** Example genome views of ARID1A occupancies on mesodermal TF EOMES, cardiac TFs HAND2, GATA4, GATA5, ISL1 and TBX3. **g** List of essential cardiogenic and cardiac functional genes occupied by ARID1A. **h** ChIP-qPCR validation of cardiogenic gene promoters occupied by ARID1A. **i** Example genome views of ARID1A occupancies on neurogenic genes. **j** List of essential neurogenic and neural marker genes occupied by ARID1A. **k** ChIP-qPCR validation of neurogenic gene promoters occupied by ARID1A. All bars in ChIP-QPCR data are shown as mean ± SD. n=3, *p < 0.05 (anti-ARID1A vs. IgG, an unpaired two-tailed t-test).

### ARID1A associates with different transcriptional factors

We predicted potential co-TFs locating within ARID1A binding regions based on their motif sequence information achieved from ARID1A ChIP-seq (Fig. 7a). TBXT (T), Repressor element-1 silencing transcription/neuron-restrictive silencer factor (REST/NRSF) and MEF2C were the top three predicted co-TFs (Fig. 7b). REST/NRSF was known to specifically repress neural gene transcription [38]. We then reanalyzed the published REST/NRSF ChIP-seq data (ENCODE: ENCSR663WAR) generated in H1 hESCs. A notable overlap was observed between REST and ARID1A occupied genes (Fig. 7c). There were total 1,113 genes co-occupied by ARID1A and REST (*p* = 9.3E-27). 28.0% (312 out of 1113) of overlapped genes were neural genes (*p* = 2.1E-57) (Fig. 7d) and 4.2% (47 out of 1113) were cardiac genes (*p*=0.06) (Fig. 7e), indicating a strong bias that ARID1A and REST tend to co-occupy more neural genes than cardiac genes. The lists of neural and cardiac genes were from an unbiased database, the Human Protein Atlas database (https://www.proteinatlas.org/). Next, integrated data analyses were performed on 40 essential neurogenic (Fig. 7f) and cardiogenic genes (Fig. 7g) showing descending differential chromatin accessibility (DCA). The purple (1^th^ column) and green (2^nd^ column) bars represented whether the gene was occupied by ARID1A and REST, respectively. The 3^rd^ column in Figs. 7f and 7g indicated signal differences of DCA during cardiac commitment. The most right two columns (4^th^ and 5^th^ column) showed relative gene expressions in WT and ARID1A^-/-^ hESC-derived cells post cardiac differentiation. Same as shown in Figs. 7d-e, REST and ARID1A co-bound much more neurogenic genes (Fig. 7f) than cardiogenic genes (Fig. 7g). REST and ARID1A co-occupied neural genes which showed various changes of chromatin accessibility, including increased, decreased, or non-changed accessibility (Fig. 7f, DCA). However, expression levels of all these neurogenic genes increased in the middle of (Figs. 5f, 5j) and after differentiation of ARID1A^-/-^ cells when compared with WT cells (Fig. 7f, 4^th^ and 5^th^ column). These results suggest that ARID1A-REST/NRSF interaction could possibly repress transcription of co-occupied neurogenic genes. As comparison, REST barely co-occupied with ARID1A on promoter regions of essential cardiogenic genes in Fig. 7g.

**Fig. 7.**
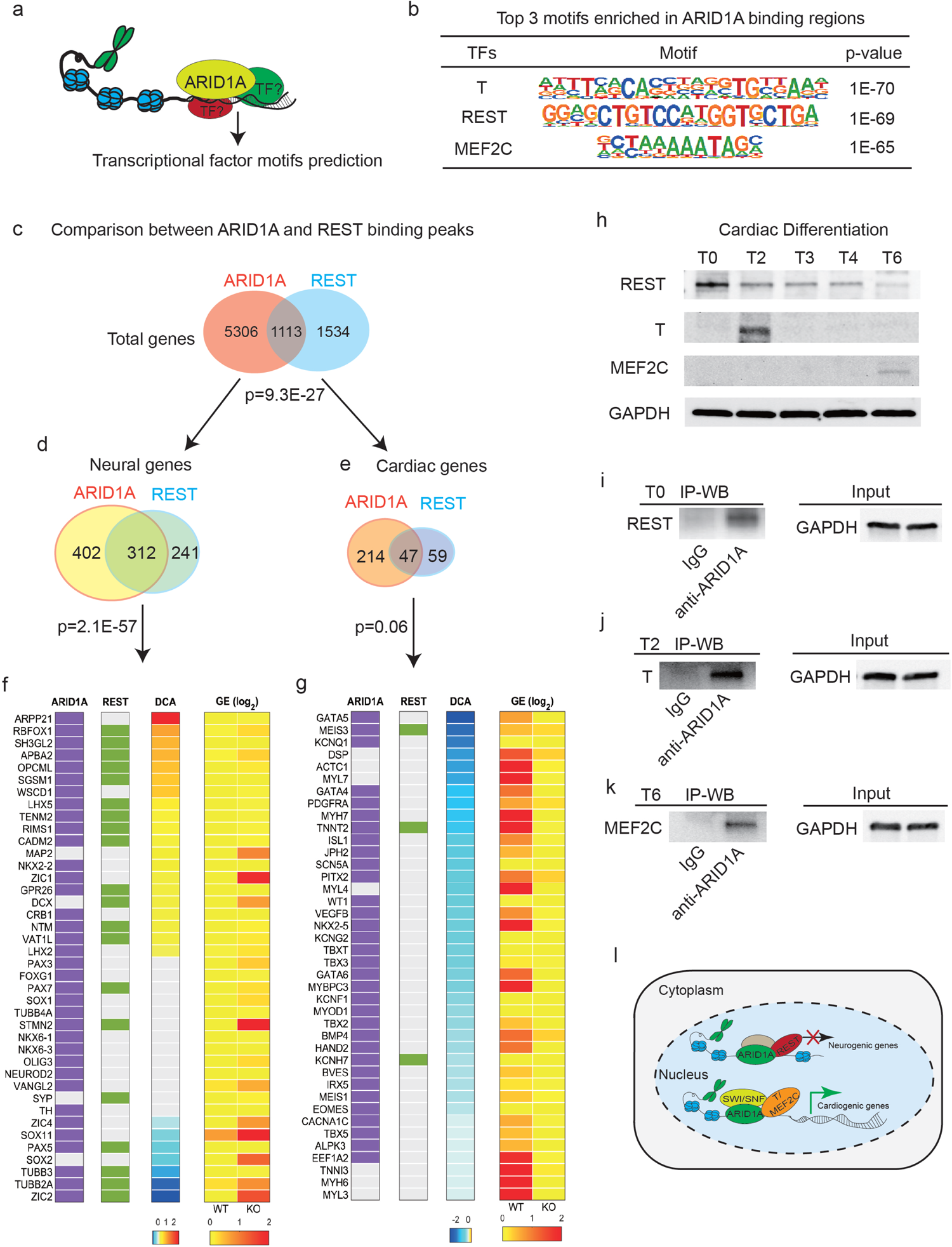
ARID1A interacts with co-factors. **a** Prediction of co-factors interacting with ARID1A according to transcriptional factor binding motifs in ARID1A ChIP-seq peaks. **b** Top three transcription factor motifs surrounding ARID1A cis-regulatory binding elements. **c** Comparison between ARID1A and REST occupied gene lists from ChIP-seq datasets. Hypergeometric model was used to calculate the significance of overlap between ARID1A and REST target genes, and p-value of enrichment of genes in both ARID1A and REST targets. **d-e** Comparison between ARID1A and REST occupied neural (**d**) and cardiac genes (**e**) from ChIP-seq datasets, indicating that REST and ARID1A prefer to co-occupy neural genes than cardiac genes. **f** Integrative analysis of ARID1A ChIP-seq (1^th^ column), REST ChIP-seq (2^nd^ column), differential chromatin accessibility (DCA) by ATAC-seq (KO vs. WT) (3^rd^ column) and average gene expression levels (GE, log2 (fold change)) from scRNA-seq (KO vs. WT, 4^th^ and 5^th^ columns). Purple boxes (1^th^ column) show the representative ARID1A occupied genes from Fig. 7d, while green boxes (2^nd^ column) show REST occupancies on those genes. **g** Same scheme design as Fig. 7f to show ARID1A and REST occupancies on representative cardiogenic genes are shown. **h** Western blotting to detect temporal protein expressions of REST, T, MEF2C during cardiac differentiation of hESCs. Protein samples were collected on day 0 (T0), 2 (T2), 3 (T3), 4 (T4) and 6 (T6). **i-k** Co-IP validation of the interaction between ARID1A and REST in undifferentiated hESCs (T0) (**i**), the interaction between ARID1A and T in hESCs at day 2 of differentiation (T2) (**j**), and the interaction between ARID1A and MEF2C in hESCs at day 6 of differentiation (T6) (**k**). **l** Schematic summary of ARID1A’s roles in human cardiac and neural commitment from hESCs.

Lastly, we verified the interactions of ARID1A with REST, T and MEF2C using co-immunoprecipitation (Co-IP) assay. Western blotting was performed to detect temporal protein levels of REST, T and MEF2C during cardiac differentiation of WT hESCs (Fig. 7h). REST expressed through all time points with gradual decrease. T showed peak expression at day 2 (T2) of mesoderm formation. MEF2C, a key cardiac TF, expressed from day 6 (T6). Anti-ARID1A antibody pulled down REST in hESCs (Fig. 7i), suggesting ARID1A deficiency in hESCs could dissociate ARID1A-REST interaction. Moreover, anti-ARID1A antibody could pull down T at day 2 (Fig. 7j) and MEF2C at day 6 (Fig. 7k) of cardiac differentiation, and vice versa (Figs. S7d-e), suggesting that ARID1A could subsequently interact with T and MEF2C to regulate chromatin accessibility during human heart development. Taken together, our results reveal the molecular mechanisms by which ARID1A regulates early human cardiac and neural development from hESCs (Fig. 7l).

## Discussion

In this study, we found ARID1A promoted human cardiac commitment via increasing chromatin accessibility on essential cardiogenic genes and suppressed neural commitment via repressing transcription of neurogenic genes. Thus, we uncovered the distinct mechanisms by ARID1A, which at chromatin remolding and transcriptional levels, to control human cardiogenesis and neurogenesis from pluripotent stem cells.

The classical function of SWI/SNF complex is to remodel chromatin state and DNA accessibility to transcription factors, hence critical for transcriptional regulation and gene expression. Arid1a (Baf250a) has been previously reported to play important role in mouse heart formation. Loss-of-Arid1a resulted in embryonic lethality with lack of primitive streak or mesoderm formation [15, 17]. Arid1a ablation in mouse second heart field (SHF) leads to trabeculation defects in the right ventricle [39]. Although previous in vivo studies revealed the critical role of Arid1a in mouse heart morphogenesis, how ARID1A drives early stage human cardiogenesis from pluripotent stem cells remains elusive. Our data demonstrate that ARID1A promotes human cardiac development from hESCs by globally increasing chromatin accessibility on essential cardiogenic genes, which include key mesoderm formation genes, cardiac specific transcriptional factors and sarcomeric genes. During cardiogenesis, temporal interaction of ARID1A with key mesoderm formation gene T and cardiac specific transcriptional factor MEF2C could very likely facilitate the sequential deposition of SWI/SNF complex on some cardiogenic genes, which regulate cardiac mesoderm specification and cardiac functionality. Although this interaction will be further studied, our data indicate that early human cardiac fate commitment from hESCs is largely dependent on the chromatin remodeling role of SWI/SNF complex.

SWI/SNF complex was also found to regulate neurodevelopment in mouse. Mice with loss of Brg [40], Baf47 (also known as Smarcb1) [41] or Baf155 (also known as Smarcc1) [42] die in pre- or peri-implantation stages. Targeted deletion of Brg1, which is an ATPase of SWI/SNF complex, in the developing mouse nervous system causes smaller brains that lack cerebellum [43]. These neurodevelopmental defects could be due to failure of neural progenitor self-renewal and differentiation [43, 44]. However, compared with the role of Brg1 in mouse neural development, our data show that loss-of-ARID1A prominently enhanced neurogenesis from hESCs. Knockout-of-ARID1A in hESCs increased neural differentiation under both targeted cardiac and neural differentiation conditions. Since the cardiac differentiation protocol was specifically modified to support mesoderm and cardiomyocyte differentiation, the robust generation of neural cells from ARID1A^-/-^ hESCs under cardiac differentiation conditions strongly suggests the master regulatory role of ARID1A in governing neurogenesis. The increased neural differentiation of ARID1A^-/-^ hESCs and under-developed brain in Brg1 KO mice indicate that ARID1A might have SWI/SNF unrelated functions in neurogenesis. This was proved by our finding that ARID1A did not uniformly affect the chromatin accessibility on neurogenic genes, instead it globally controlled the transcriptional activity of neurogenic genes. We found loss-of-ARID1A induced sporadic neural differentiation together with increased expression of multiple neurogenic genes in hESCs. Given the heterogeneity of hESCs, this observation suggests that chromatin accessibility of neurogenic genes in subpopulation(s) of undifferentiated hESCs is very likely under a relatively open status. So that loss of ARID1A could activate transcription of those genes. In fact, multiple previous studies reported that neural-associated genes in hESCs are under a bivalent condition with enhanced chromatin and transcriptional plasticity, and tend to be transcribed [45-47].

Pervious study showed that disruption of Arid1a-DNA interaction by a single mutation (Arid1aV^1068G/V1068G^) [15] in mouse caused neural tube defects including open head folds, neural tube closure defects, and heart defects such as trabeculation defects, hypoplastic myocardial walls and ventricular septal defects. Although it was not defined whether the neural tube closure defects in Arid1aV^1068G/V1068G^ mouse was due to increased neurogenesis, it was previously reported that increased number of neural stem cells led to defective neural tube closure [48-50]. Our ATAC-seq and ChIP-seq in hESCs revealed that loss-of-ARID1A globally reduced chromatin accessibility on promoters of cardiogenic genes and activated transcription of neurogenic genes. Taken together, results from Arid1aV^1068G/V1068G^ mouse model and ARID1A^-/-^ hESCs demonstrate that ARID1A plays a conserved and critical role in controlling both cardiogenesis and neurogenesis.

Our data show that ARID1A interact with REST/NRSF. During embryogenesis, REST/NRSF is widely expressed and plays a critical role in neuronal development. In pluripotent stem cells and neural progenitors, REST represses a large pool of neuron-specific genes essentail for synaptic plasticity and remodeling, including synaptic vesicle proteins, neuroreceptors and ion channels [51-53]. However, how ARID1A-REST recognizes target neurogenic genes remains elusive. Previous report in HEK293 cells found that BRG1 enhanced REST-mediated repression at some REST target genes by increasing the interaction of REST with the local chromatin at its binding sites [54]. Whether this interaction was through ARID1A was not clear. Additionally, Pax6, which is a master neuroregulatory gene, interacts with neural progenitor BAF (npBAF) complex to regulate neurogenesis [55]. Therefore, it is likely that Pax6, or other TFs, could guide ARID1A-REST to target neurogenic genes during differentiation of hESCs. Since ChIP-seq revealed ARID1A occupancy on essential cardiogenic and neurogenic genes in undifferentiated hESCs, this suggests that cardiogenesis and neurogenesis could possibly have been pre-determined by epigenetic machineries at the pluripotent stem cell stage. It also gives us a hint that ARID1A-SWI/SNF could associate with factor(s), other than REST, to repress spontaneous cardiac differentiation in hESCs. Our future study will explore this mechanism.

## Conclusion

This study uncovers the essential and opposite roles of ARID1A in governing cardiogenesis and neurogenesis from human pluripotent stem cells. Our findings reveal the distinct mechanisms by which ARID1A globally enhances chromatin remodeling on cardiogenic genes and suppresses transcription of neurogenic genes.

## Methods

### Human embryonic stem cells culture, cardiac differentiation and neural differentiation

Human embryonic stem cell (hESC) line H9 was cultured on Matrigel (BD Biosciences)-coated plates in mTesR medium [56, 57]. For cardiac differentiation, a monolayer differentiation method was used as described reported [30]. Briefly, hESCs were induced with chemically defined medium (CDM3, containing RPMI/1640, AA and BSA) as the basal medium [30] containing 6 μM CHIR99021 (Tocris Bioscience) from day 0 to day 2, 5 μM XAV-939 (Tocris Bioscience) from day 2 to day 4, then maintained in CDM3 basal medium without any chemicals for total 10-13 days. Beating cardiomyocytes were observed after differentiation for 8 to13 days. For neural differentiation, H9 hESCs were dissociated and replated into 6-well plate (coated with metrigel) and induced in N2B27 medium (25 mL DMEM/F12 medium, 1X N2, 25 mL neurobasal medium, 1X B27, 1X Glutamax, 1X NEAA and 1X Penicillin-Streptomycin). After 4 days differentiation, cells were collected for experiments.

### Vector cloning

Specific shRNA1 and shRNA2 targeting human ARID1A were cloned into the pLKO.1-TRC-puro vector (Addgene), separately. ShRNA scramble, as negative control without targeting any human genes, was also cloned into pLKO.1-TRC-puro vector.

### CRISPR/Cas-9 mediated DNA manipulation

All gRNAs were designed based on CRISPR design platform from MIT (http://crispr.mit.edu/). Full length of human ARID1A gene was knocked out by dual gRNAs in hESC H9. For each gRNA oligo, self-complementary oligos were purchased from Invitrogen. Both of gRNAs, targeting 5’ sequence and 3’ sequence of ARID1A DNA, were cloned into the pENTR-spCAS9-T2A-EGFP vector (from Dr. Yi Sheng), separately. Single gRNA, targeting and disrupting ARID1A TSS, was cloned into the LentiCRISPRv2-puro vector (Addgene) [58]. pENTR-spCAS9-T2A-EGFP-gRNAs vector were transfected into H9 hESCs using X-treme GENE 9 DNA Transfection Reagent (Roche). After 24h transfection, GFP positive cells were sorted by FACSAria II cell sorter (BD Biosciences) and replated to grow single clones in mTesR medium with ROCK inhibitor Y-27632 (Tocris Bioscience). Single H9 clones were picked out and replated into 48-well plate for further expansion. Genomic DNAs of single H9 clones were extracted by the DNeasy Blood & Tissue kit (QIAGEN) kit. Null ARID1A hESC clones were verified by PCR using different primer sets. For ARID1A TSS disruption, H9 cells were infected with lentiCRISPRv2-puro-ARID1A gRNA virus and puromycin was added to select drug resistant clones. LntiCRISPRv2-puro-ARID1A gRNA vector-mediated genomic DNA editing on human ARID1A promoter was detected by Surveyor® Mutation Detection Kit for Standard Gel Electrophoresis (Integrated DNA Technologies, Inc.).

### Surveyor assay

Genomic DNA editing was detected using Surveyor® Mutation Detection Kit for Standard Gel Electrophoresis (Integrated DNA Technologies) according to manufacturer’s instructions. Briefly, genomic DNA was extracted by DNeasy Blood & Tissue Kit (QIAGEN). PCR fragments were purified by GeneJET PCR Purification Kit (K0701). After that, PCR fragments were annealing in 1X prime star buffer (Takara) and Surveyor enzyme was added to digest the annealed DNA fragments at 42 °C for 1h. Gel Electrophoresis on agarose gel was used to detect the digestion.

### Lentivirus package and transduction

The lentiviral vector lentiCRISPRv2-puro was transfected into the HEK293T cells (ATCC) along with packaging plasmids including psPAX2 and pMD2.G (from Dr. Guang Hu in NIH) using the X-treme GENE 9 transfection reagent (Roche). Viral supernatant was collected and cellular debris was removed by syringe filtering (0.45 μm pore size; Millipore). Human H9 cells cultured in mTesR medium were incubated with virus media for 4h, followed with fresh mTesR medium culture for overnight. Same infection was repeated after 24 hr. Puromycin was added to select puromycin-resistant H9 clones after virus infection for 48h.

### PCR screening of ARID1A^-/-^ clones

HESC clones were picked and expanded in mTesR medium. Genomic DNAs from each hESCs clone were purified by DNeasy Blood & Tissue Kits (Qiagen). PCRs with different primer sets were conducted by using DreamTaq Green PCR Master Mix (Thermo Scientific) or PrimeSTAR DNA Polymerase kit (Takara). PCR products were run on agarose gel to visualize the fragment size. For all primers, please see the Table S4.

### RNA extraction and RT-Quantitative PCR

Total RNAs were extracted by miRNeasy mini kit (Qiagen) or RNeasy Mini Kit (Qiagen). High-Capacity RNA-to-cDNA™ Kit (Applied Biosystems) was used for cDNA synthesis. RT-Quantitative PCR (RT-qPCR) was performed on a 7900HT Fast Real-Time PCR System (Applied Biosystems) with Fast SYBR Green Master Mix (Applied Biosystems). The RT-qPCR results were normalized to internal control GAPDH or beta-ACTIN using the 2^−ΔΔCt^ method as previously described [59]. RT-qPCR data were presented as mean ± S.D. from at least three independent experiments.

### Western Blotting

Cells were lysed and proteins were purified by Complete™ Lysis-M EDTA-free kit (Roche, 04719964001). Protein samples were run on Mini-PROTEAN® TGX™ Precast Gels (Bio-Rad) and transferred to PVDF membranes by a wet transfer apparatus (Bio-Rad). After blotting with 5% non-fat milk for 30 min at room temperature, the membranes were incubated in TBXT buffer containing 5% non-fat milk and primary antibodies at 4°C overnight. On the next day, membranes were incubated in TBXT buffer containing 5% non-fat milk and horseradish peroxidase (HRP)-conjugated anti-rabbit or anti-mouse IgG (Cell signaling tech.) and detected by using ECL Western Blotting Substrate (Pierce).

### Co-immunoprecipitation (Co-IP)

Cell proteins were extracted by Pierce™ Classic Magnetic IP/Co-IP Kit (Thermo Scientifi, 88804) or Complete™ Lysis-M EDTA-free kit (Roche, 04719964001) according to the manuals. Protein Co-immunoprecipitation (Co-IP) was performed by using Pierce™ Classic Magnetic IP/Co-IP Kit as well. Protein samples were run on Mini-PROTEAN TGX Precast Gels (Bio-Rad) and probed with appropriate antibodies by Western blotting.

### Chromatin Immunoprecipitation-qPCR (ChIP-qPCR)

H9 hESCs were cultured in P10 plate in mTesRm medium. ChIP was carried out in undifferentiated hESCs according to manuals of truChIP™ Chromatin Shearing Kit (Covaris, PN 520154) and EZ-Magna ChIP™ A/G Chromatin Immunoprecipitation Kit (Millipore, 17-10086). Briefly, hESCs were fixed with methanol-free formaldehyde provided by truChIP™ Chromatin Shearing Kit (Covaris, PN 520154). Sonication of cell lysis was done by a ME220 Focused-ultrasonicator using truChIP Chromatin Shearing Tissue Kit (Covaris). Chromatin immunoprecipitation (ChIP) was performed by EZ Magna ChIP™ A/G Chromatin Immunoprecipitation Kit (Millipore). Human ARID1A antibody (Millipore: (04-080) Anti-BAF250a/ARID1a Antibody, clone PSG3) was used to pull down genomic DNAs. Normal mouse/rabbit IgG or RNA Polymerase II antibodies (provided by Millipore ChIP kit) were used as negative or positive control, respectively. ChIP-qPCR signals were calculated as fold enrichment of 1% Input or non-specific antibody (isotype IgG antibodies) signals with at less three technical triplicates. Each specific antibody ChIP sample was normalized to its isotype IgG antibody-ChIP-signals obtained in the same sample. Standard deviations (SD) were calculated from technical triplicates and represented as error bars.

### ChIP-sequencing (ChIP-seq)

H9 hESCs were cultured in P10 plate in mTesRm medium. ChIP was carried out in undifferentiated hESCs according to manuals of truChIP™ Chromatin Shearing Kit (Covaris, PN 520154) and EZ-Magna ChIP™ A/G Chromatin Immunoprecipitation Kit (Millipore, 17-10086). Briefly, hESCs were fixed with methanol-free formaldehyde provided by truChIP™ Chromatin Shearing Kit (Covaris, PN 520154). Chromatin of cell lysis was sheared using truChIP™ Chromatin Shearing Kit according its manual by ME220 Focused-ultrasonicator (Covaris). The sheared chromatin was then incubated with anti-ARID1A antibody and purified by using EZ-Magna ChIP™ A/G Chromatin Immunoprecipitation Kit (Millipore, 17-10086). Chromatin DNAs quality was assessed by 2100 Bioanalyzer (Agilent Technologies) and sequenced in the Center for Medical Genomics at Indiana University School of Medicine.

### Single cell 3’ RNA sequencing (scRNA-seq)

H9 hESCs and differentiated hESCs were dissociated by Corning™ 0.25% Trypsin (Corning™ 25053CI) to single cells. Single cell 3’ RNA-seq experiments were conducted using the Chromium single cell system (10x Genomics, Inc) and Illumina sequencers at the Center of Molecular Genetics (CMG) of Indiana University School of Medicine. Cell suspension was first inspected on the Countess II FL (Thermo Fisher Scientific) and under microscope for cell number, cell viability, and cell size. Depending on the quality of the initial cell suspension, the single cell preparation included centrifugation, re-suspension, and filtration to remove cell debris, dead cells and cell aggregates. Single cell capture and library preparation were carried out according to the Chromium Single cell 3’ Reagent kits V2 User Guide (10X Genomics PN-120267, PN-1000009, PN-120262). Appropriate number of cells were loaded on a multiple-channel micro-fluidics chip of the Chromium Single Cell Instrument (10x Genomics) with a targeted cell recovery of 10,000. Single cell gel beads in emulsion containing barcoded oligonucleotides and reverse transcriptase reagents were generated with the v2 single cell reagent kit (10X Genomics). Following cell capture and cell lysis, cDNA was synthesized and amplified. Illumina sequencing libraries were then prepared with the amplified cDNA. The resulting libraries were assessed with an Agilent TapeStation or Bioanalyzer 2100. The final libraries of the undifferentiated samples were sequenced using a custom program on Illumina NextSeq 500/550; and the libraries of the differentiated samples were sequenced on Illumina NovaSeq 6000. 26 bp of cell barcode and UMI sequences, and 91 or 98 bp RNA reads were generated with Illumina NextSeq500/550 or NovaSeq 6000.

### Analysis of scRNA-seq data

CellRanger 2.1.0 (http://support.10xgenomics.com/) was utilized to process the raw sequence data generated. Briefly, CellRanger used bcl2fastq (https://support.illumina.com/) to demultiplex raw base sequence calls generated from the sequencer into sample-specific FASTQ files. The FASTQ files were then aligned to the human reference genome GRCh38 with RNA-seq aligner STAR. The aligned reads were traced back to individual cells and the gene expression level of individual genes were quantified based on the number of UMIs (unique molecular indices) detected in each cell. The filtered gene-cell barcode matrices generated with CellRanger were used for further analysis with the R package Seurat (version 2.3.1) [28] with Rstudio version 1.1.453 and R version 3.5.1. Quality control (QC) of the data was implemented as the first step in our analysis. We first filtered out genes that were detected in less than five cells and cells with less than 200 genes. To further exclude low-quality cells in downstream analysis we used the function isOutlier from R package scater [29] together with visual inspection of the distributions of number of genes, UMIs, and mitochondrial gene content. Cells with extremely high or low number of detected genes/UMIs were excluded. In addition, cells with high percentage of mitochondrial reads were also filtered out. After removing likely multiplets and low-quality cells, the gene expression levels for each cell were normalized with the NormalizeData function in Seurat. To reduce variations sourced from different number of UMIs and mitochondrial gene expression, we used the ScaleData function to linearly regress out these variations. Highly variable genes were identified (x.low.cutoff=0.0125, x.high.cutoff=4, y.cutoff=0.5).

To integrate the single cell data from undifferentiated or differentiated WT and ARID1A^-/-^ samples, we applied the canonical correlation analysis (CCA) in Seurat. We chose the top 1500 variable genes from each sample to calculate the correlation components (CCs) and used the function MetageneBicorPlot to determine the optimal number of CCs. We retained the cells whose expression profiles could be explained with at least 50% by the CCs using CalcVarExpRatio and SubsetData. The CCA subspaces were then aligned with AlighSubspace using the number of CCs determined. We employed FindClusters for shared nearest neighbor (SNN) graph-based clustering. The clusters were visualized with t-distributed stochastic neighbor embedding (t-SNE) by running dimensionality reduction with RunTSNE and TSNEPlot. The FindConservedMarkers function was subsequently used to identify canonical cell type marker genes that are conserved across WT and knockout cells. To compare average gene expression within the same cluster between WT and knockout cells, we applied AverageExpression. R packages ggplot2 (ISBN 978-3-319-24277-4) and ggrepel (https://github.com/slowkow/ggrepel) were used to plot the average gene expression. Violin plots (VlnPlot) and feature plots (FeaturePlot) were used to visualize specific gene expressions across clusters and different sample conditions.

### Assay for Transposase Accessible Chromatin with high-throughput sequencing (ATAC-seq)

WT and ARID1A^-/-^ hESCs were differentiated for 4 days, then cells were dissociated by Trypsin-EDTA (0.25%). Cells were washed by 1X PBS and re-suspended in cold PBS according to ATAC-seq protocol [60]. Briefly, collected cells were lysed in cold lysis buffer (10 mM Tris-HCl, pH 7.4, 10 mM NaCl, 3 mM MgCl2 and 0.1% IGEPAL CA-630) and the nuclei were pelleted and resuspended in Tn5 enzyme and transposase buffer (Illumina Nextera® DNA library preparation kit, cat# FC-121-1030). The Nextera libraries were amplified using the Nextera® PCR master mix and KAPA biosystems HiFi hotstart readymix (cat # NC0295239) successively. AMPure XP beads (Beckman Coulter cat# A63881) were used to purify the transposed DNA and the amplified PCR products. All libraries were sequenced on a 100 cycle paired-end run on an Illumina NOVAseq instrument. The resulting ATAC-seq libraries were sequenced on Illumina NovaSeq 6000 at CMG of Indiana University School of Medicine and paired-end 50 bp reads were generated. Illumina adapter sequences and low-quality base calls were trimmed off the paired-end reads with Trim Galore v0.4.3 (http://www.bioinformatics.babraham.ac.uk/projects/trim_galore/). The resulting high-quality reads were aligned to the human reference genome hg38 using bowtie2 (version 2.3.2) [61] with parameters “-X 2000 --no-mixed --no-discordant”. Duplicate reads were discarded with Picard (https://broadinstitute.github.io/picard/). Reads mapped to mitochondrial DNA together with low mapping quality reads (MAPQ<10) were excluded from further analysis. ATAC-seq was conducted in Genomic Core at Indiana University.

### Integrated analysis of ATAC-seq and ChIP-seq results

Bowtie2 [61] was used to align sequencing reads on the human genome (hg38) for both ChIP-seq and ATAC-seq. Low quality score (Q30) reads and redundant reads due to PCR bias were filtered by Samtools [62, 63] and Picard MarkDuplicates that were not considered for further analysis. There were two technical replicates for ATAC-seq for both WT and ARID1A^-/-^. Open chromatin regions for each sample were identified first from peak calling of individual ATAC-seq using MACS2 [64] with broad peak option and cutoff of q < 0.1. We then constructed a final set of unique regions for differential analysis by merging overlapped open chromatin regions recognized from different samples/conditions. The read abundance of each open region was counted for each individual sample, respectively, by using pyDNase [65]. A software, edgeR [66, 67], was employed to perform differential accessibility analysis on the read counts of each chromatin region for WT and ARID1A^-/-^ samples. Differentially accessible open chromatin regions between WT and ARID1A^-/-^ were determined by specific cutoffs, FDR-adjusted p-value less than 0.01 and amplitude of fold change (in log scale with base 2) larger than 0.5. ARID1A binding sites were determined by model-based ChIP-seq data analysis using MACS2 [64]. Peak calling of uniquely mapped ARID1A ChIP-seq reads was performed by comparing with input ChIP-seq. All binding peaks were recognized if their p-values less than 0.01 after Benjamini-Hochberg multiple-test correction. We examined consensus sequences significantly enriched in ARID1A binding peaks. The motif enrichment analysis was performed by using the HOMER (version 4.9.1) [68] command “findMotifsGenome.pl” with the parameter “-size given”. The open chromatin regions detected by ATAC-seq or ARID1A binding sites identified by ARD1A ChIP-seq were linked to specific genes, if they locate either within upstream 10kb from TSS, or 5’UTR, or exon/intron (gene body) of the genes. The UCSC genome browser custom track was made to visualize differences of chromatin accessibility between WT and ARID1A^-/-^ for selected regions. The read numbers were normalized by total numbers of high-quality reads in each sample, then multiplied by ten millions. The normalized averaged counts for replicates under the same condition were presented in the figures.

### Flow cytometry

Flow cytometry was performed according to our previous publication [69]. Briefly, cells were harvested and dissociated by 0.25% trypsin-EDTA for 5 -10 min at 37°C. The dissociated single cells were fixed in 4% PFA (diluted with 16% Paraformaldehyde (formaldehyde) aqueous solution) for 10 mins at room temperature and washed 3 times with 1X PBS. Cells were incubated in blocking PBS buffer containing 2% goat serum and 0.1% saponin. Then cells were incubated in blocking buffer with primary antibody for 1h at 37°C, following with secondary antibody staining for 1 hr at 37°C. Flow cytometry analysis was carried out with Accuri C6 flow cytometer (Becton Dickinson), BD LSRII cytometer (Becton Dickinson) and Attune NxT Flow Cytometer (Thermo Fisher Scientific). Data were analyzed by FlowJo (Treestar).

### Immunofluorescence (IF)

For immunocytochemistry, cells were fixed with 4% PFA (diluted from 16% Paraformaldehyde aqueous solution) for 10 min at room temperature. After washing with 1X PBS, cells were blocked for 1 h with 1X PBS blocking buffer containing 2% goat serum (or 5% BSA) and 0.1% saponin. Staining with primary antibodies diluted with blocking buffer was performed for overnight at -4°C. Staining with secondary antibodies were performed on next day, following with nucleus staining with DAPI. Leica DM6B image system was used for imaging.

### Functional enrichment analysis

The functional enrichment analysis, including gene ontology (GO), process networks, signaling pathways, and genes interaction networks, was performed on gene sets selected by using Metacore software (Clarivate Analytics). We chose the cutoff *p* < 0.05 to determine the functions significantly over-represented in genes of our interest.

### Quantification and Statistical Analysis

For RT-QPCR and flow cytometry data, comparisons between two groups (KO versus WT) were conducted using an unpaired two-tailed *t*-test. All data were presented as mean ± S.D. from at least three independent experiments. Differences with *P* values less than 0.05 were considered significant. A nonparametric test, Wilcoxon signed-rank test, was used to compare the gene expression difference between WT and ARID1A^-/-^ detected by scRNA-seq. The Fisher’s test was adopted to determine the statistical significance of difference of ratios of cell numbers between WT and ARID1A^-/-^. We used hypergeometric model to calculate the significance of overlap between ARID1A and REST target genes, and *p*-value of enrichment of cardiac or neural genes in both ARID1A and REST targets.

## Acknowledgments

We thank Matthew David Durbin from Indiana University School of Medicine for reading the manuscript.

## Funding

This work was supported by 2014 NIH Director’s New Innovator Award (1DP2HL127727-01) and 2019 American Heart Association (AHA) Transformative Project Award (19TPA34850038) to L.Y, NIH Indiana University Simon Cancer Center Support Grant (P30CA082709) to J.W, AHA Postdoctoral Fellowship (17POST33460140) to J. L.

## Author Contributions

JL and LY initiated and designed studies. JL performed most experiments. LH performed western blot. XZ and YW worked on ATAC-seq experiments, YS and W.S provided guidance for CRISPR/Cas9 technology, YL provided guidance about scRNA-seq. HG analyzed scRNA-seq data. SL and JW performed bioinformatics analysis on ChIP-seq and ATAC-seq, and did integrative analysis to include other published datasets. JL, JW and LY interpreted data/results and wrote the manuscript. LY supervised the whole research and provided guidance. All authors had access to the final manuscript and approved the submission of the article.

## Competing interests

The authors declare no competing interests.

## Figure legends

**Figure S1:**
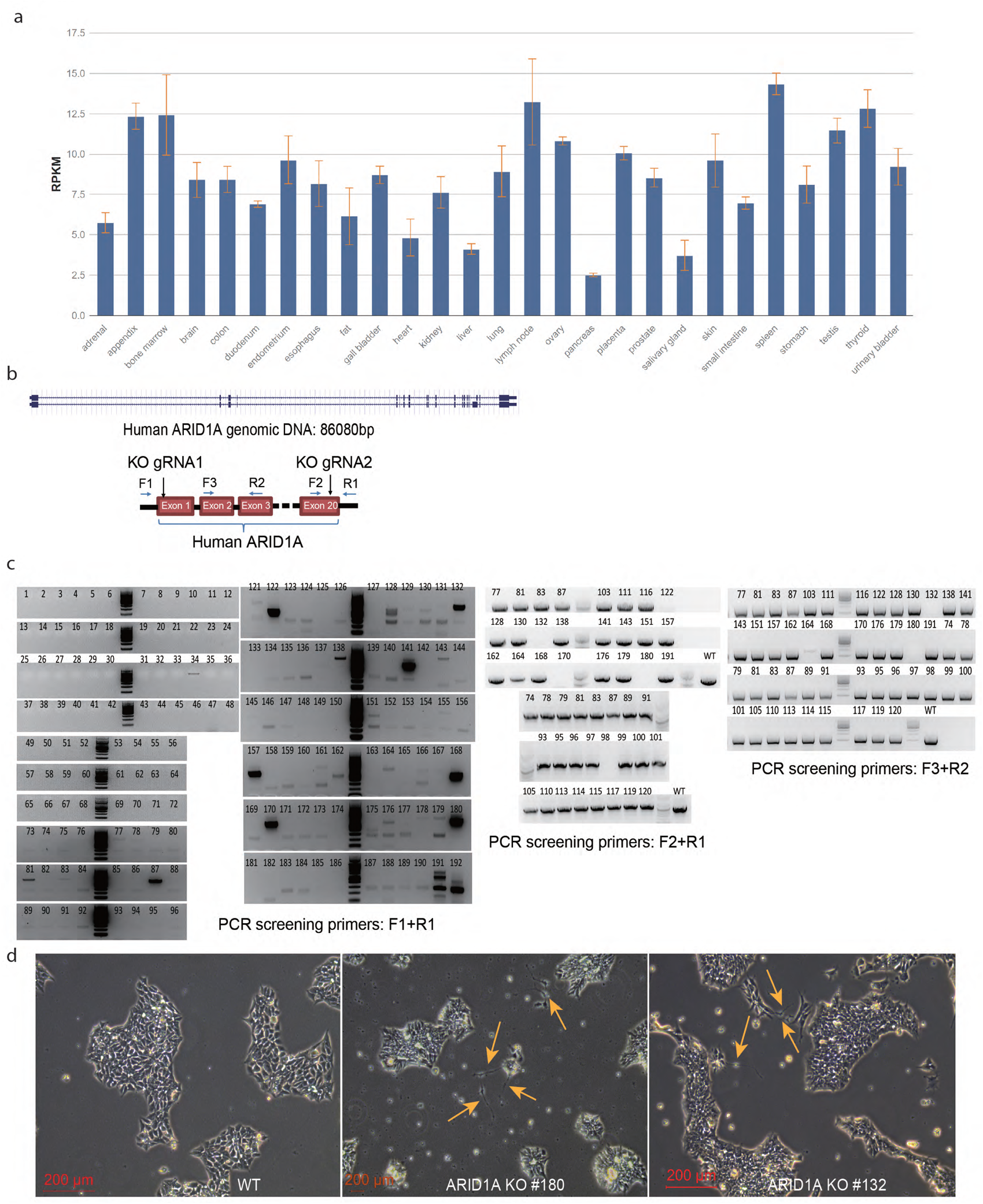
Establishment of ARID1A null H9 hESC line. **a** ARID1A expression in different human samples from NCBI database. **b** ARID1A was completely knocked out from hESC H9 cell line by using dual gRNAs mediated by CRISPR/Cas9 technology. **c** Identification of ARID1A knockout (KO) hESC clones by using PCR. **d** WT and ARID1A KO hESCs cultured in mTesR medium. WT hESCs show normal stem cell morphology. ARID1A KO hESCs give rise to small spontaneous differentiated cell clusters (yellow arrows).

**Figure S2:**
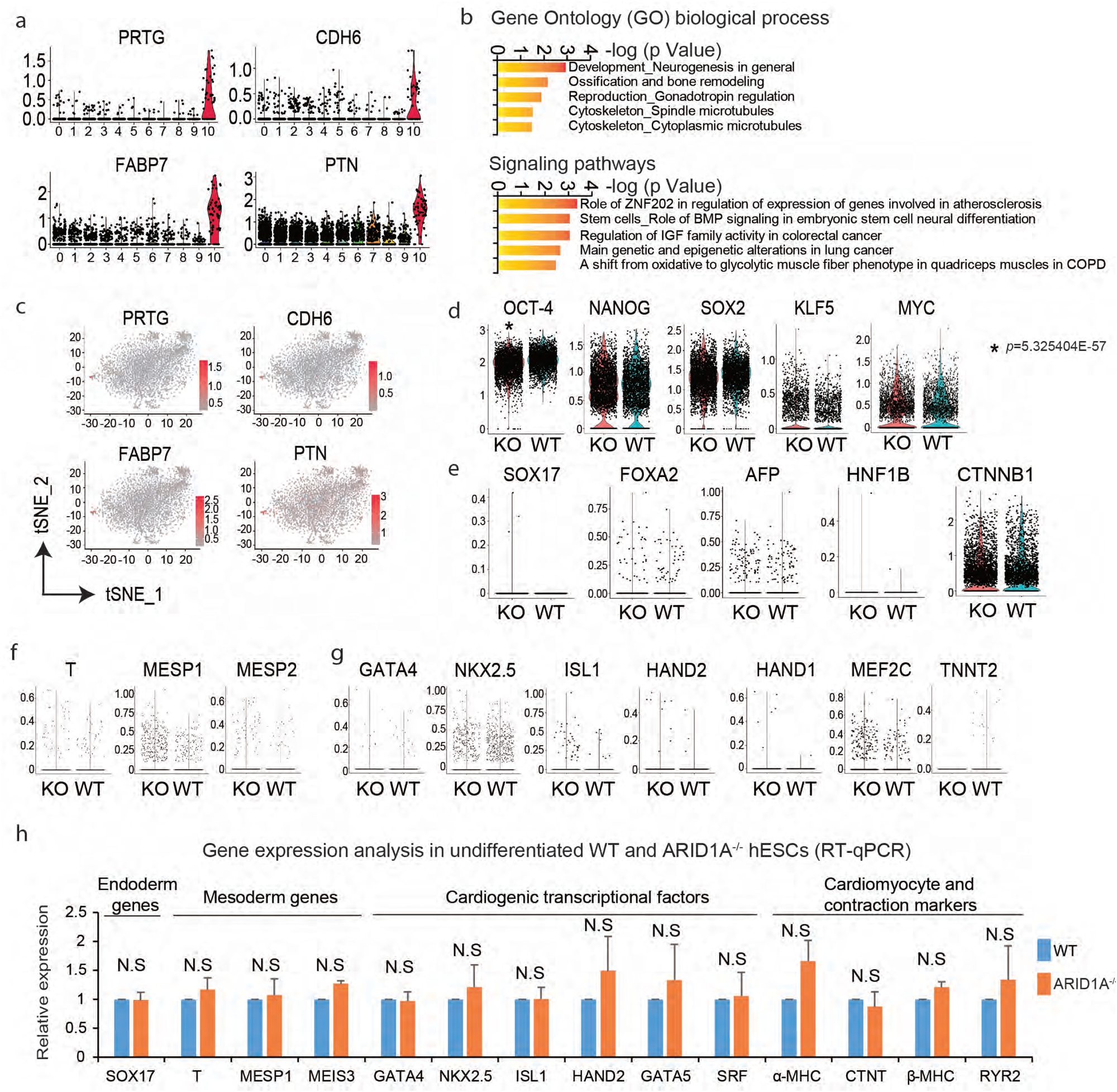
ScRNAseq analysis of undifferentiated WT and ARID1A^-/-^ hESCs. **a** Violin plots analysis of neural markers in each cluster. **b** GO biological process (top) and signaling pathways (bottom) analysis of upregulated genes (KO vs. WT) in the cluster 10. Analysis was performed by Metacore software. **c** Feature plots of neural markers in integrative WT and KO scRNA-seq data. **d** Violin plots analysis of pluripotency markers in WT and KO hESCs. * p=5.325404E-57 (Nonparametric Wilcoxon signed-rank test). **e** Violin plots analysis of endodermal markers in WT and KO hESCs. All p values were larger than 0.05 (KO vs. WT), by Nonparametric Wilcoxon signed-rank test. **f** Violin plots analysis of early mesodermal markers in WT and KO hESCs. All p values were larger than 0.05 (KO vs. WT), by Nonparametric Wilcoxon signed-rank test. **g** Violin plots analysis of cardiac and cardiomyocyte markers in WT and KO hESCs. All p values were larger than 0.05 (KO vs. WT), by Nonparametric Wilcoxon signed-rank test. **h** RT-qPCR verification showing gene expression in undifferentiated WT and ARID1A^-/-^ hESCs. N.S, no significance (ARID1A^-/-^ vs. WT), by unpaired two-tailed t-test.

**Figure S3:**
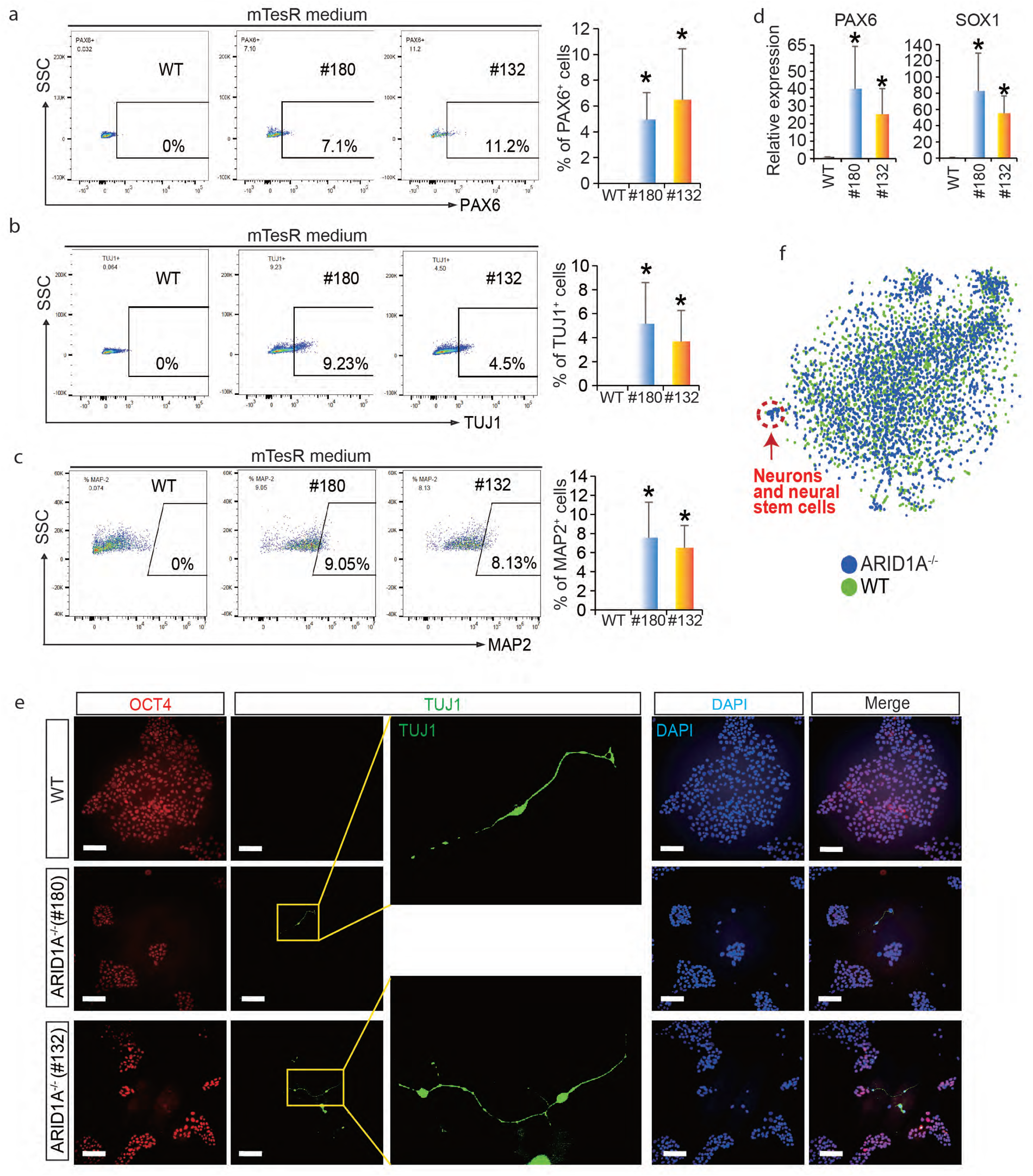
Identity of neural cells from ARID1A^-/-^ hESCs. **a** Quantification of the ratio of PAX6^+^ cells derived from undifferentiated WT and KO hESCs (#180, #132) by flow cytometry. **b** Quantification of the ratio of TUJ1^+^ cells derived from undifferentiated WT and KO hESCs (#180, #132) by flow cytometry. **c** Quantification of the ratio of MAP2^+^ cells derived from undifferentiated WT and KO hESCs (#180, #132) by flow cytometry. **d** RT-qPCR showing the expression levels of PAX6 and SOX1 from undifferentiated WT and KO hESCs (#180, #132). **e** Immunofluorescence analysis of pluripotency marker OCT4 and neuron marker TUJ1 in WT and KO hESCs (#180, #132). Scale bar, 100 µm. **f** Interactive analysis of scRNA-seq datasets from undifferentiated WT and KO hESCs. All bars are shown as mean ± SD. n=3, *p < 0.05 (an unpaired two-tailed t-test with Welch’s correction).

**Figure S4:**
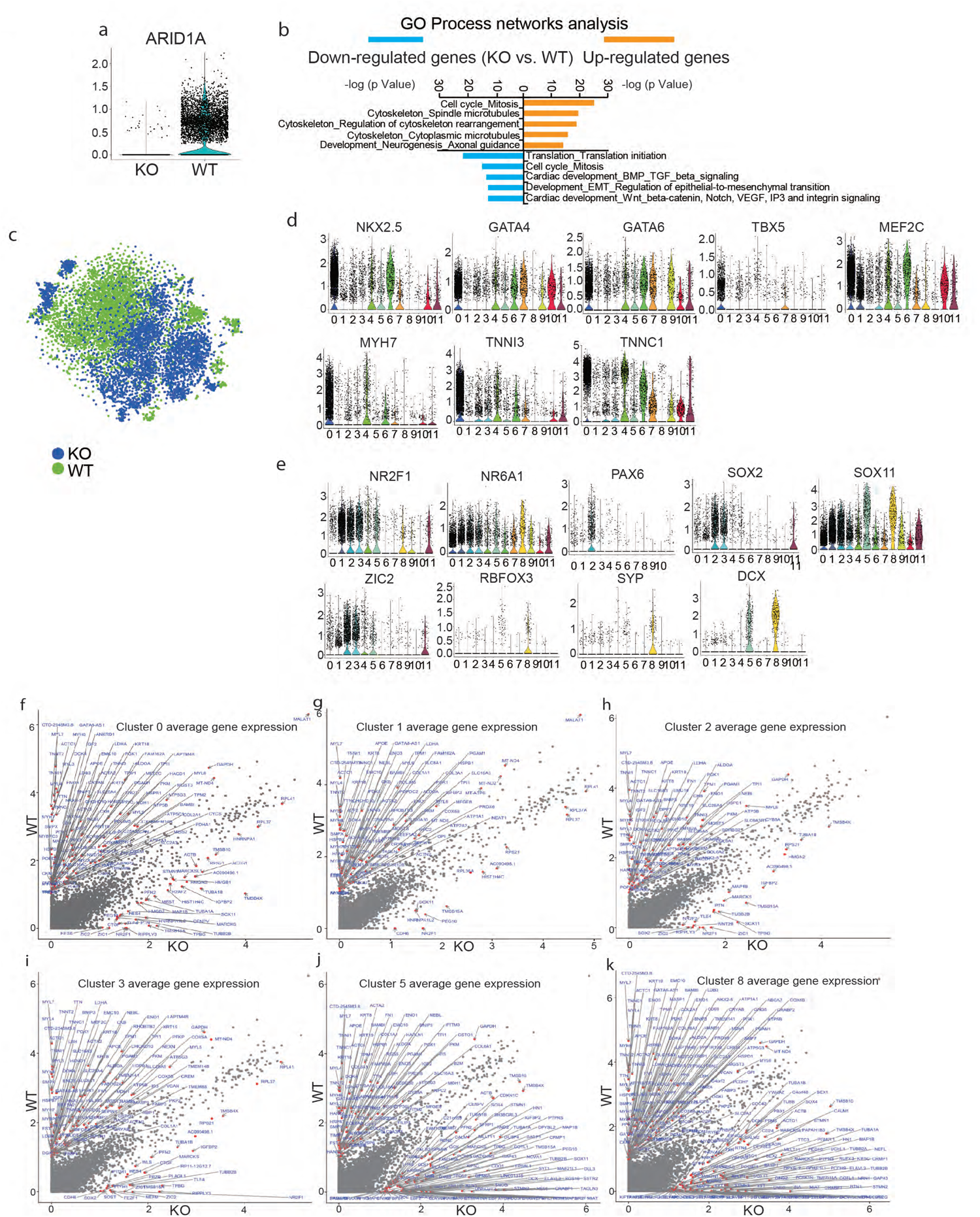
ScRNA-seq reveals loss-of-ARID1A represses cardiac but promotes neural differentiation from hESCs. **a** ARID1A expressions in WT and KO hESC-derived cells by scRNA-seq. **b** GO process networks analysis of down-regulated genes and up-regulated genes (KO vs. WT) by Metacore software. **c** Integrative analysis of scRNA-seq data from differentiated WT and KO cells. Blue blots showed the KO cells and green blots showed the WT cells. **d** Violin plots analysis of cardiogenic gene expressions in each cluster shown with integrative WT and KO scRNA-seq dataset. **e** Violin plots analysis of neurogenic gene expressions in each cluster shown with integrative WT and KO scRNA-seq dataset. **f-k** Differential gene expression analyses in cluster 0 (**f**), cluster 1 (**g**), cluster 2 (**h**), cluster 3 (**i**), cluster 5 (**j**) and cluster 8 (**k**).

**Figure S5:**
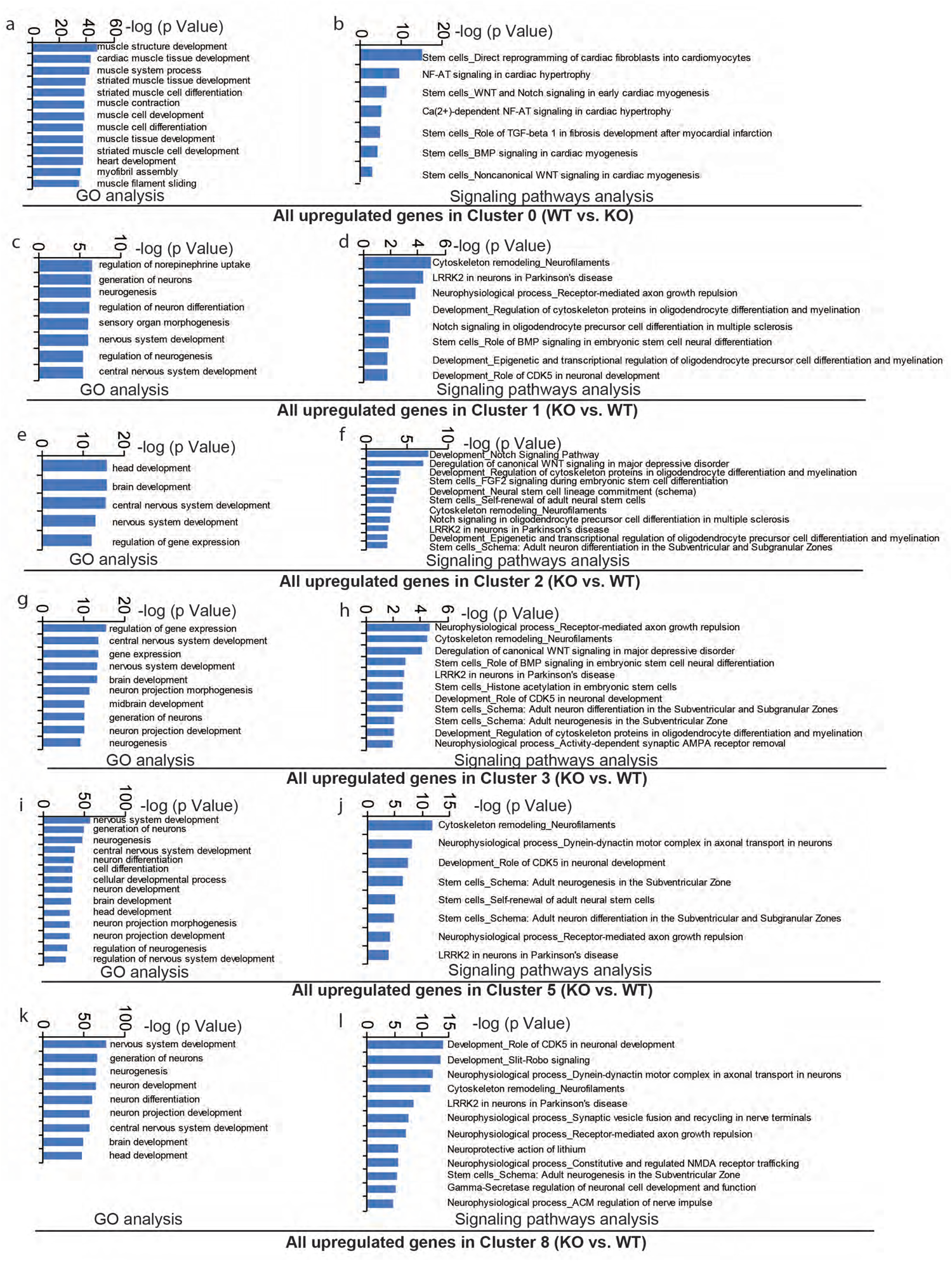
Functional enrichment analysis. **a-b** GO biological process (left) and signaling pathways (right) analysis of all upregulated genes (WT vs. KO) in cluster 0. **c-l** GO biological process (left) and signaling pathways (right) analysis of all upregulated genes (KO vs. WT) in Cluster (**c-d**) 1, Cluster 2 (**e-f**), Cluster 3 (**g-h**), Cluster 5 (**i-j**) and Cluster 8 (**k-l**).

**Figure S6:**
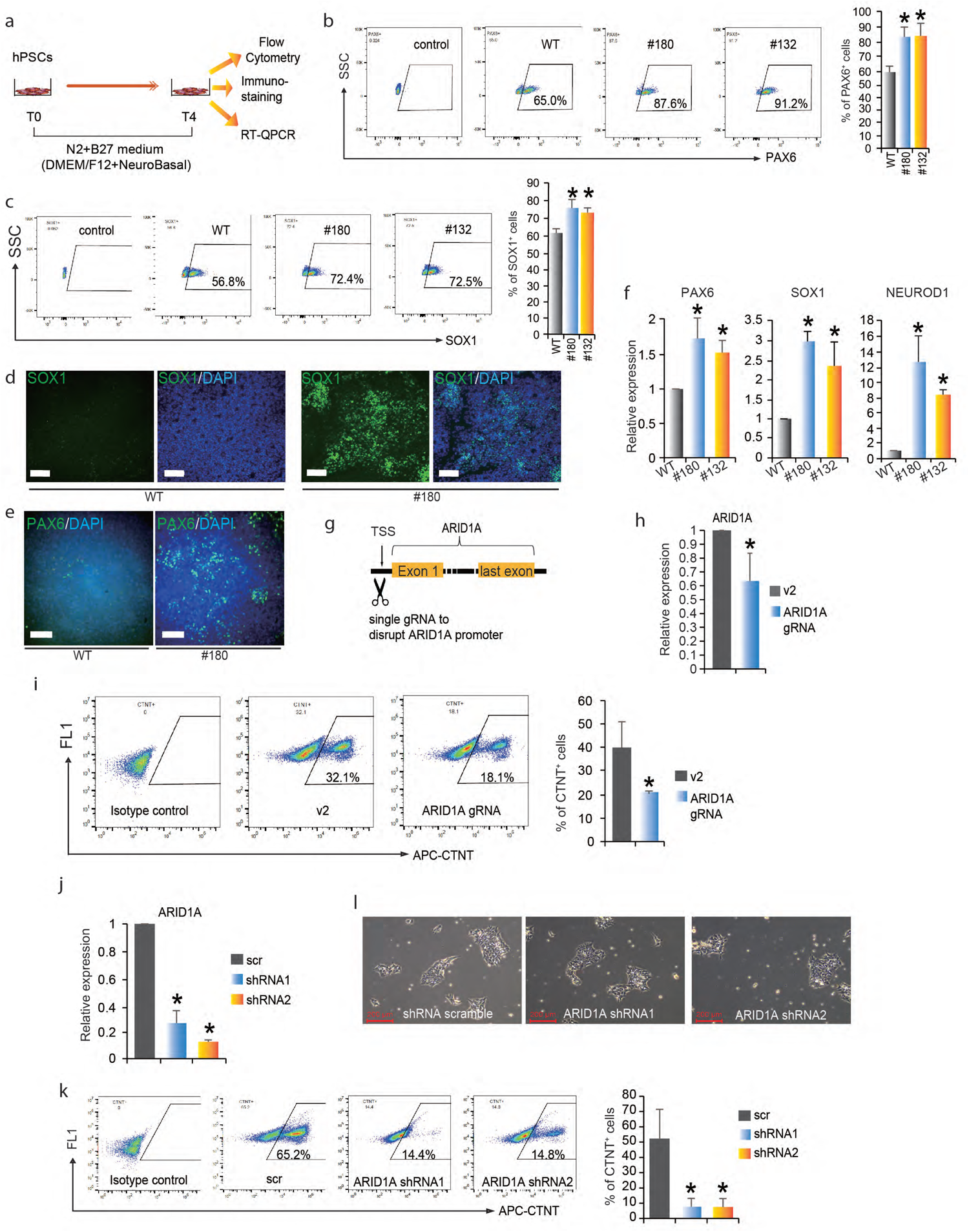
Loss-of-ARID1A promotes neural differentiation under neural differentiation conditions and knockdown of ARID1A promotes cardiac differentiation for hESCs. **a** Strategy for neural differentiation of WT and ARID1A^-/-^ hESCs. **b-c** Flow cytometry data showing increased percentages of PAX6^+^ (**b**) and SOX1^+^ (**c**) cells derived from ARID1A KO hESCs (#180, #132) post specific neural differentiation when compared to that of WT hESCs. **d-e** Loss-of-ARID1A (#180) increased the differentiation of SOX1^+^ (**d**) and PAX6^+^ (**e**) cells after neural differentiation than WT hESCs by immunostaining. Scale bar, 100 µm. **f** QRT-PCR analysis of PAX6, SOX1 and NEUROD1 mRNA expression levels post neural differentiation. **g** Human ARID1A promoter was disrupted by single gRNA (ARID1A gRNA) targeting TSS. **h** ARID1A mRNA expression was decreased by ARID1A TSS-gRNA in hESCs. **i** Flow cytometry data showing knockdown of ARID1A decreased percentage of CTNT^+^ cells compared to that of WT, after 10 days of cardiac differentiation. **j** ARID1A expression levels were decreased by two shRNAs in H9 hESCs. **k** ARID1A shRNAs decreased percentage of CTNT^+^ cells after 10 days of cardiac differentiation. **l** Both of WT and ARID1A^knockdown^ hESCs were cultured in mTesR medium, showing normal stem cell morphology without any visible differentiated cells. Scale bar, 200 µm. All bars are shown as mean ± SD. n=3, *p < 0.05 (an unpaired two-tailed t-test).

**Figure S7:**
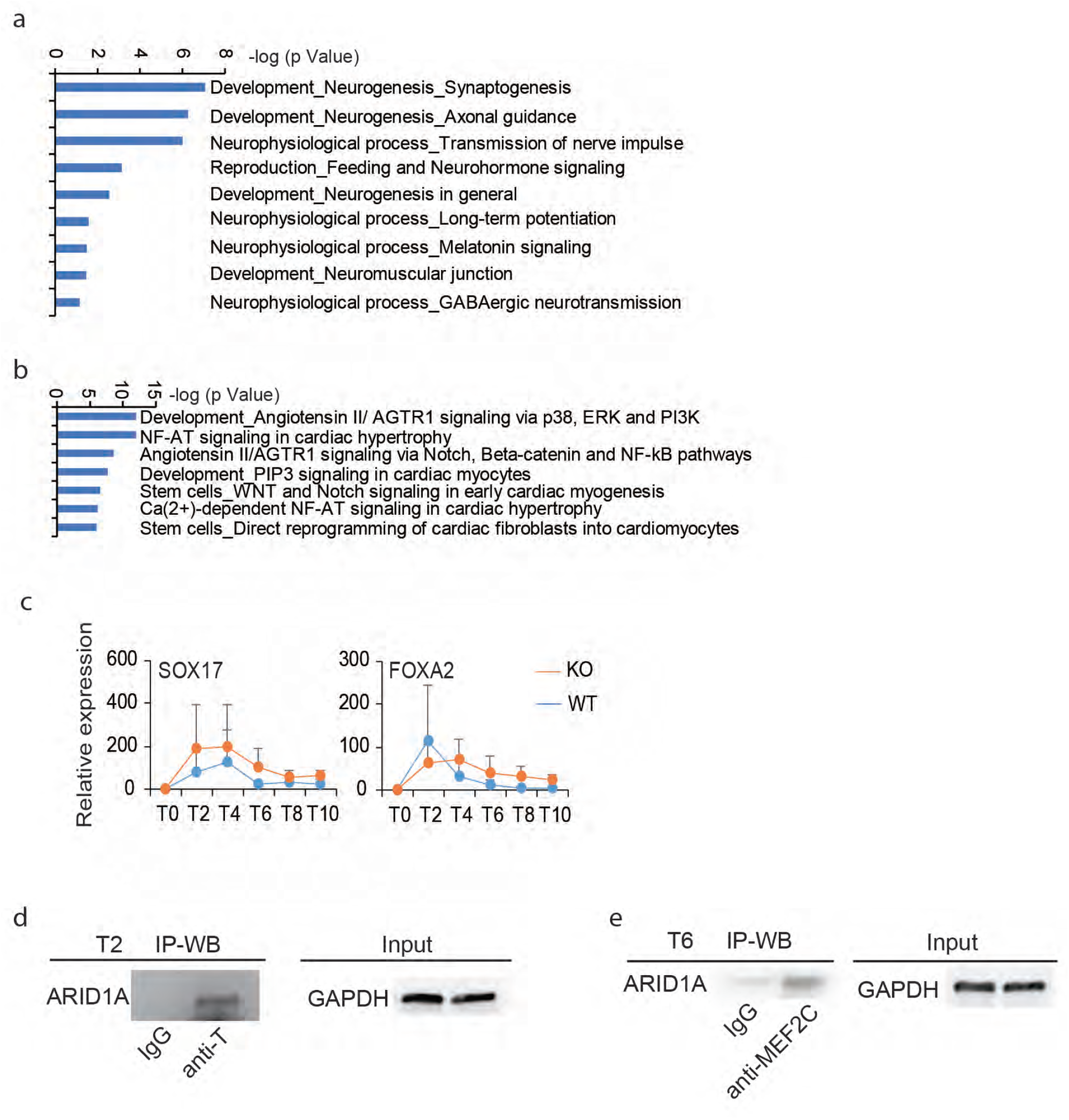
ARID1A affects chromatin accessibility and interacts with other transcriptional factors. **a** Process networks analysis of genes with increased chromatin accessibilities (WT vs. KO) by Metacore. **b** Signaling pathways analysis of genes with decreased chromatin accessibilities (WT vs. KO) by Metacore. **c** Temporal mRNA expression profiles of endodermal markers SOX17 and FOXA2 during cardiac differentiation. WT and KO hESCs were differentiated from day 0 (T0) to day 10 (T10), and RNAs were collected every 2 days. All bars are shown as mean ± SD. n=3. Unpaired two-tailed t-test with Welch’s correction. **d** Co-IP validation of interaction of T and ARID1A by using anti-T antibody to pull down ARID1A after 2 days (T2) of cardiac differentiation. **e** Co-IP validation of interaction of MEF2C and ARID1A by using anti-MEF2C antibody to pull down ARID1A after 6 days (T6) of cardiac differentiation.

## Notes

### Competing Interest Statement

The authors have declared no competing interest.

